# Brain function in language and associated networks in non- or minimally verbal children

**DOI:** 10.1101/2025.11.10.686908

**Authors:** Guillermo Montaña-Valverde, Annika Linke, Dominika Slušná, Jordi Muchart-López, Antoni Rodríguez-Fornells, Gustavo Deco, Wolfram Hinzen

## Abstract

Language is universal in humans, develops robustly in infancy, and is rarely absent entirely in aphasia following strokes. Non- or minimally verbal children, in whom language has not developed by school-age in either production or comprehension, thus provide an unparalleled window into the human brain when it is deprived of language function. Yet insights from functional MRI are absent. Here we report results from a first study of intrinsic connectivity in 9 non- or minimally verbal children with autism spectrum disorder (nvASD) scanned under sedation with propofol along with 8 typically developing children (scanned awake), using resting-state (rs) fMRI. We targeted both functional (FC) and anatomically constrained, generative effective connectivity (GEC) in the speech and language networks, the insula and associated networks, and the hippocampus. Identical analyses were applied to an independent rsMRI dataset of healthy adults scanned both under propofol and while awake, to evaluate sedation confounds. NvASDs compared to their neurotypical peers showed a widespread pattern of hypoconnectivity in auditory speech perception, frontotemporal, and semantic processing regions, which extended further to the insula, and the hippocampus. GEC results selectively replicated these patterns, which correlated with autism diagnostic observation schedule (ADOS) behavioral scores within the nvASD group. This hypoconnectivity pattern extended neither to the adults scanned under propofol, nor to the visual cortex used as control region in nvASD, suggesting that this pattern of results could not be explained by sedation. Together, this first evidence from intrinsic connectivity reveals a broad pattern of underconnectivity across key cognitive networks, which provides a neural correlate for the significant breakdown of language-related cognitive functions in this population.

## Introduction

In neurotypical infants, language develops universally and reliably on a biologically timed path. While production of connected speech does not develop before the second year, speech perception evolves from before birth, while production of vowels, consonants, and syllables emerge during the first year.^1^ Even sophisticated semantic comprehension long predates the first birthday.^2–4^ At the brain level, structural asymmetries in cortical language regions, well-established in adults, are evident in preterm newborns,^5^ and the acoustic radiation is present as early as one month.^6^ Auditory cortex responds to speech in an adult-similar fashion from three months,^7–10^ when functional connectivity in speech networks is mature as well.^11^

These findings illustrate a neural infrastructure of language present from the beginning of a neurotypical human life, and thus contribute to the mystery of why a considerable number of children do not develop language by school-age. This clinical phenotype of ‘absent speech’^12^ is not rare, signalling a failure of a key developmental outcome across a large range of neurogenetic conditions. In a case like perisylvian polymicrogyria, brain malformations affecting the cortex around the sylvian fissure provide an anatomical basis for this phenotype,^13^ which can be accompanied by severely compromised receptive language as well.^14,15^ The arcuate fasciculus, a key white matter tract connecting speech-motor and auditory regions, has also often been documented to be undetectable in this population.^16^ No such pertinent anatomical irregularities are seen in the instance of non- or minimally verbal autism (nvASD), which forms a sizable 30% of the autism spectrum. Also missing in nvASD are gross motor deficits, which form a key part of the clinical presentations of other syndromes that can involve absent speech, such as Worster-Drought,^17^ Landau-Kleffner,^18^ or Cri du chat syndromes.^19^ In many syndromes, absent speech also goes along with relatively more preserved comprehension, particularly in movement disorders involving primary apraxia of speech, e.g. Angelman,^20^ Coffin Siris,^21^ or Phelan-McDermid syndromes.^22^ Again, no such asymmetry is seen in nvASD, where speech comprehension has been reported not to exceed production^23^, which would effectively replicate a pattern seen in language abilities in ASD at large^24^ (a subset of 25% of children identified as minimally verbal demonstrated higher receptive compared to productive language).^107^ In those children with nvASD that can produce or learn single words, apprehension of meaning has been found to be highly anomalous.^25^

NvASD forms the centre of this study and was here defined as comprising a global failure of language development by school age, covering both production and comprehension. This crucially entails an absence of evidence of language abilities in any other physical medium, such as writing or a formal Sign language, the existence of which would immediately contradict the term ‘nonverbal’, which should never be confused with ‘nonspeaking’.^26^ So defined, this profile does not rule out non-linguistic *communication* abilities, such as those based on pictograms,^27^ which can be present and are trainable in many children with nvASD. Absence of language in nvASD is accompanied by intellectual disability in a large majority, and by broad nonverbal cognitive impairment such as impairments in perceptual categorization or understanding nonverbal representations (e.g., pictures).^23,28,29^ While human cognition in the absence of language development remains a virtually uncharted territory,^30^ nvASD makes it pertinent to investigate language dysfunction in the context of a broader neurocognitive breakdown and, in particular, not to focus it around the notion of *speech*. Speech is the phonetic articulation of language (how language *sounds*), not language itself, as comprised by meaning and structure. While even brain-structural insights in this population are scarce,^31^ neurofunctional studies of language at any level are largely absent, as they are impeded by the difficulty of conducting neuroimaging studies in awake children at this end of the autism spectrum, particularly in paradigms involving tasks.^32^ Ethical restrictions on sedating neurotypical controls along with clinical cases further contribute to a profound lack of insights in this domain.

Studying spontaneous fluctuations in the fMRI BOLD signal during the ‘resting state’ reveals networks generally consistent with those revealed under cognitive tasks, including in the case of the language network and in children.^33–38^ Moreover, there is striking evidence for preservation of intrinsic functional networks under anaesthesia,^39,40^ or even in some instances of the vegetative state.^41^ On the other hand, notable differences in functional connectivity (FC) due to sedation have also been reported, notably in higher-order associative networks such as the default mode network (DMN), while primary sensory networks are reported as unaffected under sedation.^42^ Recently, functional connectivity in the form of correlations between time series of different brain regions has been supplemented by the more mechanistic framework of generative effective connectivity (GEC), which improves FC by weighting the existing anatomical connectivity and thus approximating the asymmetrical causal impact of one region on another.^43,44^

Here we reasoned that this evidence warrants a first study of intrinsic connectivity in lightly sedated children with nvASD, which were specifically selected here to form a maximally homogeneous group in terms of their language profile as characterized above. We thus performed a case study of 9 subjects with nvASD using rs fMRI, using an analysis scheme involving both FC as a baseline and GEC, targeting primary auditory speech perception regions at first, radiating out from there to the extended language network including the semantic network and the hippocampus. Within-group correlations to behavioral measures, and an independent dataset of young adults scanned both awake and sedated, were used to guard against sedation confounds. Specifically, starting from primary auditory cortex, we included frontotemporal perisylvian regions (inferior frontal (IFG) and superior/middle temporal gyri (S/MTG)). The semantic network was included because any normal act of language use involves meaning,^45^ which as such takes us beyond the confines of the classical neurologically identified language regions and in fact comprises large swaths of association cortex,^46^ overlapping strongly with the medial frontoparietal DMN,^47^ a key vulnerability across major mental disorders including ASD.^48^ The semantic network also includes the anterior insula, which we targeted separately and jointly and the ‘salience network’ due to their key status across major mental disorders including ASD,^49^ and their role in controlling higher-order functions related to the DMN.^50^ The hippocampus was included as it has been argued to form part of the language network,^51,52^ and to play a key role in language learning,^53^ semantic memory,^54,55^ pronoun processing,^56^ and the verbal compression and recall of narrative information.^57^

Extensive individual heterogeneity in the autism spectrum extends to available evidence on intrinsic functional brain connectivity. This evidence has been largely confined to the higher-functioning regions of the (verbal) autism spectrum and exhibits a complex pattern of over- and under-connectivities. A well-attested pattern is less segregation (marked by overconnectivity within functional networks) with more diffuse connectivity between networks (marked by underconnectivity) and inter-hemispherically.^58^ In task-based language studies with fMRI, a pattern of increased FC within the language network jointly with decreased FC between language and regions outside of the language network (‘global’ connectivity) is discernible,^59^ potentially interpretable as reflecting heightened processing of lower-level linguistic features such as phonemes or individual words, along with decreased integrative processing across networks. Such integration is critical for higher-order language functions such as complex syntax, discourse and narratives. Intrinsic FC in the resting state, however, does not consistently replicate this pattern.^59^ One rare study of lower-functioning (but verbal) children with ASD demonstrates a pattern of underconnectivity within the default mode network and the ventral visual stream, relative to a higher-functioning ASD group, which in turn showed a pattern of predominantly reduced network segregation in relation to neurotypical controls.^60^

While it is difficult to make predictions for nvASD based on this evidence, the uniform absence of language was our critical criterion of recruitment, which made us expect less heterogeneity in this subgroup. Given the absence of connected speech, the above pattern of overconnectivity in lower-level speech perception regions was certainly not expected, while we did expect the pattern of underconnectivity at the global or between-network level. This would also be consistent with the empirical fact that a broad breakdown of higher-order cognition is seen in nvASD, and also with the fact that language mediates key cognitive functions in development,^61^ including the social-cognitive functions so critical to the ASD cognitive phenotype.^62,63^

## Materials and methods

### nvASD dataset

This study included nine non- or minimally verbal school-aged children and adolescents diagnosed with nvASD (see Table I), recruited from special education schools in Barcelona, Spain. Recruitment criteria included: (a) an ASD diagnosis reported by parents or the clinical centre, confirmed during recruitment using both the Autism Diagnosis Observation Schedule (ADOS)^64^ and the Autism Diagnosis Interview-Revised (ADI-R)^65^; (b) an absence of phrase-level functional speech. This study was approved by the corresponding institutional review board (CEIC Fundació Sant Joan de Déu; PIC-99-17). Written informed consent was obtained from legal guardians of all participants. Given the nonverbal nature of our participants, consent was parental in all cases, following established protocols. Special attention was given to monitoring non-verbal assent, and efforts were made to minimize discomfort, including observing participants willingness to engage, and ensuring the presence of parents. These measures were in accordance with the best practices from Catalunya’s leading pediatric hospital. To provide a comparison to typical development, we included a control group of eight typically developing (TD) children. These children had no personal or family history of ASD or other neurological or psychiatric conditions and were recruited to match the ASD group in age, handedness and sex. This study was in accordance with the 2000 Helsinki declaration and its later amendments.^66^

**Table 1.**
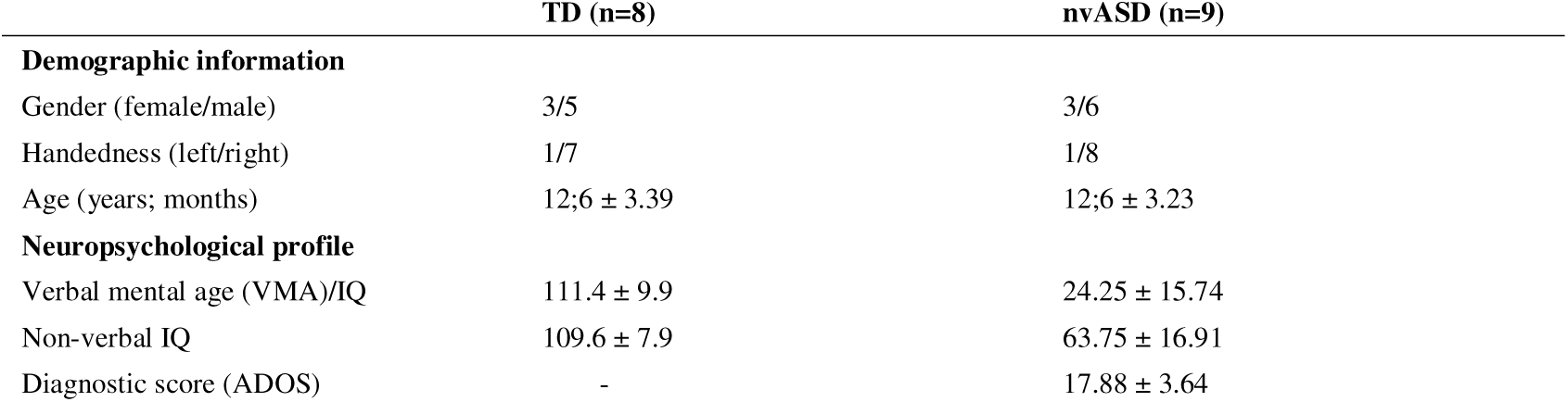
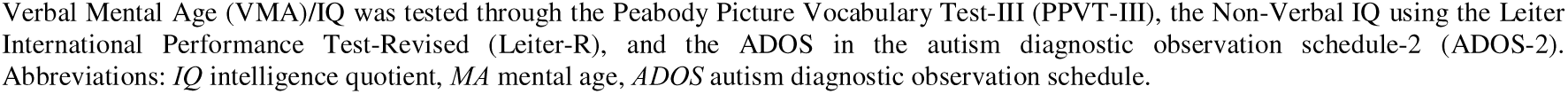
Demographic and neuropsychological participant profile.

|  | TD (n=8) | nvASD (n=9) |
| --- | --- | --- |
| **Demographic information** |  |  |
| Gender (female/male) | 3/5 | 3/6 |
| Handedness (left/right) | 1/7 | 1/8 |
| Age (years; months) | 12;6 ± 3.39 | 12;6 ± 3.23 |
| **Neuropsychological profile** |  |  |
| Verbal mental age (VMA)/IQ | 111.4 ± 9.9 | 24.25 ± 15.74 |
| Non-verbal IQ | 109.6 ± 7.9 | 63.75 ± 16.91 |
| Diagnostic score (ADOS) | - | 17.88 ± 3.64 |

Verbal Mental Age (VMA)/IQ was tested through the Peabody Picture Vocabulary Test-III (PPVT-III), the Non-Verbal IQ using the Leiter International Performance Test-Revised (Leiter-R), and the ADOS in the autism diagnostic observation schedule-2 (ADOS-2). Abbreviations: *IQ* intelligence quotient, *MA* mental age, *ADOS* autism diagnostic observation schedule.

#### Sedation procedure

Due to practical difficulties in conducting behavioural assessment in individuals with nvASD and to guard against head movements, sedation was used in this group following standard procedures in the San Joan de Déu hospital. Anaesthesia was induced via a mask with sevoflurane prior to the transition to an intravenous-based anaesthetic with propofol. Propofol dosage for induction was 1 mg/kg and after the circulation of the drug in the patient was established, a dose of 8-10 mg/kg/h was administered. Dosage gradually decreased to 6 mg/kg/h until the end of the procedure. The duration of sedation using both sevoflurane and propofol is very small allowing for fast recovery.

### Propofol healthy control comparison dataset

To test the impact of propofol sedation on functional connectivity in the neurotypical brain, we used data from a previously published study made publicly available on the OpenNeuro data repository (doi:10.18112/openneuro.ds003171.v2.0.0).^68,69^ Initially, this included data from 17 healthy subjects (mean age = 24 ± 5 years; 4 females, 13 males), but two were excluded due to a sedative period under 50 timepoints, leaving a final sample of 15 subjects, right-handed, native speakers who showed no history of neurological disorders. Participants underwent an fMRI scan while listening to an audio clip and then at rest, across four sequential levels of consciousness: awake, light sedation, deep sedation, and recovery. For our study, we only utilized the measurements from the awake and deep sedation resting states. Deep sedation in the OpenNeuro dataset was obtained through a propofol infusion with a concentration of 0.9 µg/ml, following an initial infusion of 0.6 µg/ml during the light sedation state. Oxygen levels were adjusted to maintain SpO2 above 96%. Participants were considered deeply sedated (Ramsey level 5) when they stopped responding to verbal commands and were unable to converse. At this point, the participant was maintained at the deep sedation level while retaining spontaneous cardiovascular function and ventilation. Figure 1 illustrates the process from raw data to FC and GEC matrices, as detailed in subsequent sections.

**Figure 1.**
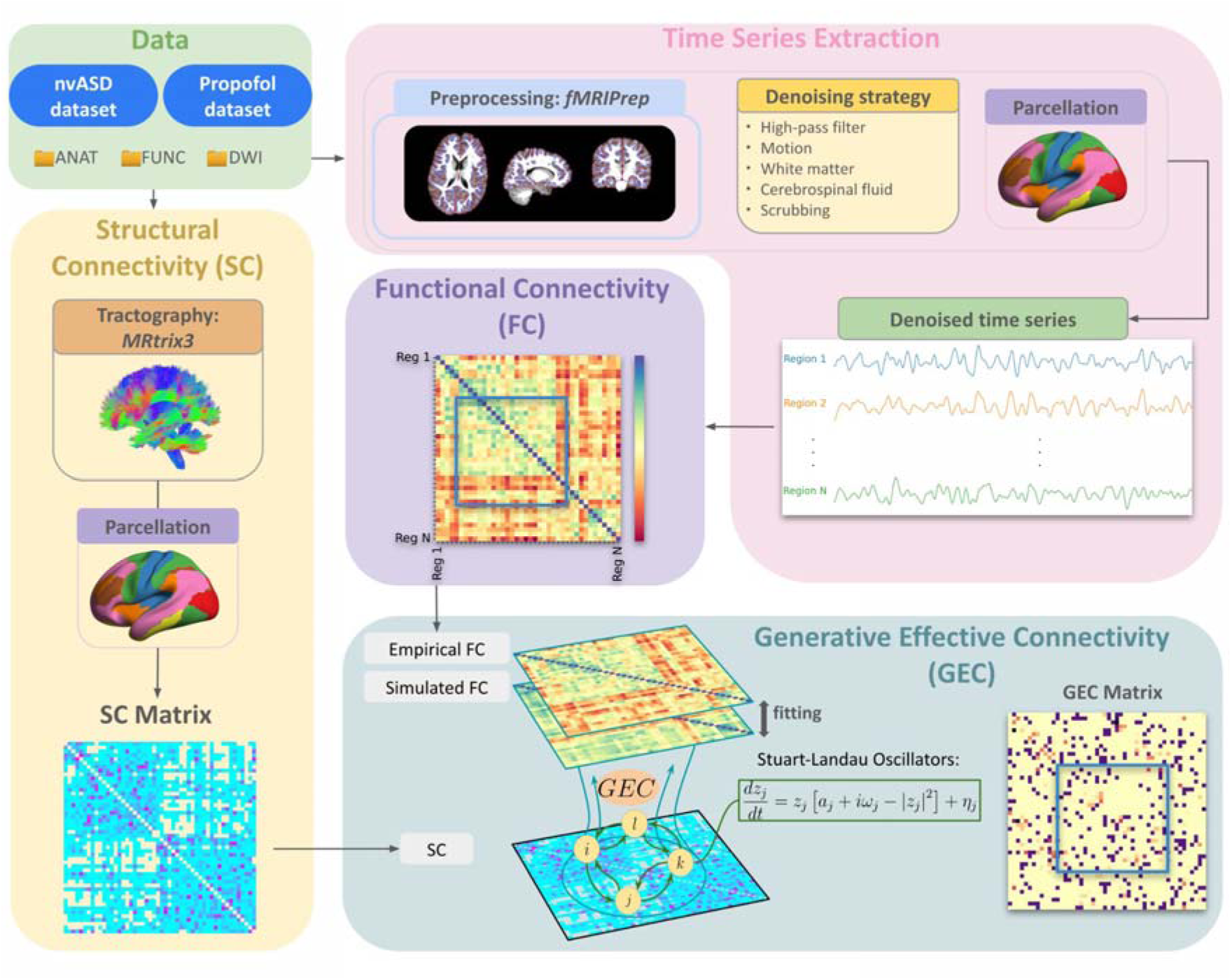
Methodological procedure. Time series were extracted from denoised, preprocessed functional MRI data using the Schaefer atlas for brain parcellation. Functional connectivity (FC) was then obtained for each subject by calculating the z-scored Pearson correlation between each pair of time series across all regions and networks of interest. This FC matrix, together with the structural connectivity (SC) matrix derived from diffusion weighted imaging (DWI) tractography, was used to compute the anatomically constrained, generative effective connectivity (GEC). FC and EC involving every set of regions of interest (blue squares) were compared between groups in each dataset.

### MRI acquisition

#### nvASD dataset

Both high-resolution T1-weighted structural images and resting state fMRI scans were acquired on a Philips Ingenia 3T scanner with a 64-channel head coil at the Sant Joan de Déu Hospital in Barcelona. The high-resolution T1-weighted structural images were obtained using a Magnetization Prepared - Rapid acquisition Gradient Echo (MPRAGE) sequence with the following parameters: repetition time (TR) = 9.899 msec, echo time (TE) = 4.6 msec, flip angle = 8°, slice thickness = 1 mm, in-plane resolution = 1mm, 180 transverse slices, and a matrix size of 240 x 240.

Resting state scans were acquired using a blood-oxygen-level-dependent (BOLD) echo planar T2*-weighted gradient echo sequence with the following parameters: TR = 2000 msec, TE = 30 msec, flip angle = 70°, acquisition matrix = 68 x 68, in-plane resolution = 3.5 mm, slice-thickness = 3.5 mm, 32 transverse slices aligned to the plane intersecting the anterior and posterior commissures. The functional run consisted of 266 volumes, with a total duration of 8 minutes and 52 seconds.

Diffusion MRI data were acquired with the following sequence parameters: Field-of-view = 230×230 mm; voxel-size = 2.05×2.05mm^2^; repetition time (TR) = 10.1s; echo time (TE) = 102ms; flip angle = 90□; number-of-slices = 64; slice-thickness = 2.1mm; number of averages = 1; acceleration factor = 2; number of shells = 2; b-values = 625, and 1250 s/mm^2^; number of diffusion gradient directions = 36; number of b0 (i.e., b-value=0) images = 3, plus 1 b0 with reverse phase to correct for spatial distortions.

#### Propofol control dataset

Participants were provided with noise cancelling headphones (Sensimetrics, S14; www.sens.com). Imaging was performed on a 3T Siemens Tim Trio system with a 32-channel coil and all subjects underwent a rs fMRI scans while awake and during different levels of propofol sedation (33 slices, voxel size: 3mm³ isotropic, inter-slice gap of 25%, TR=2000 ms, TE = 30 ms, matrix size = 64×64, FA = 75°). The awake and deep sedation states consisted of 250 (SD=12.0) and 200 (SD=73.2) volumes, respectively. Anatomical scans were also obtained using a T1-weighted 3D Magnetization Prepared - Rapid Gradient Echo (MPRAGE) sequence (voxel size: 1mm³ isotropic, TR = 2.3, TE = 4.25 ms, matrix size = 240×256×192, FA = 9°).

Diffusion MRI data were acquired following sequence parameters: Field-of-view = 192×192 mm; voxel-size = 2×2×2mm; repetition time (TR) = 9.6s; echo time (TE) = 77ms; flip angle = 90□; number-of-slices = 77; slice-thickness = 2mm; number of averages = 1; acceleration factor = 3; b-values = 700s/mm2.

### Image pre-processing

The fMRI dataset was organised in the Brain Imaging Structure (BIDS) standard so as to be passed to fMRIPrep.^70^ Detailed steps and functions are provided in the Supplementary material.

### Regressing confounds and parcellation

To reduce potential confounding effects in the fMRI data, several regressors were included in analyses. First, we applied a high-pass filter to remove low-frequency signals that can be introduced by physiological and scanner noise resources. Next, to reduce the effects of head motion, we regressed out the six rigid-body motion parameters which have been previously demonstrated that can introduce bias in group comparisons.^71^ These motion parameters include the transition on the three axes (x, y, z) and the respective rotation (a, {3, y), which are estimated relative to a reference image.^72^

Additionally, to minimize the impact of non-neuronal BOLD signal fluctuations, which are unlikely to reflect anatomical activity,^73^ we regressed out signals from white matter and cerebrospinal fluid. These signals as well as the motion parameters were expanded using the first temporal derivatives and their quadratic terms to capture potential non-linear effects of these noise sources.^74^ Furthermore, we removed high-motion segments in which the framewise displacement (FD) exceeds 0.5 mm, a process known as scrubbing.^71^ FD measures the movement of the head from one volume to the next, helping to identify and exclude volumes with excessive motion. Finally, we included the standardized DVARS, defined as the root mean squared intensity difference between two consecutive volumes, and applied a threshold of 3 to identify and exclude volumes with excessive signal intensity changes. These regression steps were based on a denoising strategy proposed by Wang *et al.*^75^ which aims to improve the quality of fMRI connectivity studies.

Finally, this regression procedure was applied to the corresponding preprocessed NifTI images through a brain atlas for denoising. In this study a custom atlas was built by the combination of the 200 parcellation Schaefer atlas together with the hippocampus and the amygdala from the Yale Brain Atlas.^76,77^ This atlas was used to assess both network and region level analysis. Each denoised time series is then saved as the main output for the following analysis.

### Seed-based FC analysis

Recent research shows that linear methods are useful for describing brain connectivity;^78^ In this study, seed-based FC analyses were used to evaluate the temporal correlations between each ROI’s resting state time course for each pair of regions. The Pearson’s correlation coefficient was calculated for each participant and z-transformed to improve normality and obtain an approximate normal distribution for statistical analysis.

### GEC analysis - The Hopf Model

The Hopf model emulates the dynamics emerging from the mutual interactions between brain areas, considered to be interconnected on the basis of anatomical structural connectivity.^43^ In other words, it combines brain structure and activity dynamics to explore and explain the underlying mechanisms of brain connectivity. In this study, we utilized a Stuart-Landau oscillator model (first introduced by Matthews and Strogatz^79^) to represent the local dynamics of each brain area described by the normal form of a supercritical Hopf Bifurcation (Supplementary Fig. 1). This is the canonical model for studying the transition from noisy, asynchronous, damped oscillations, to oscillatory, synchronous, self-sustained oscillations.^80^ The interaction between different Hopf oscillators using brain network architecture have been shown to successfully replicate features of brain dynamics observed in EEG,^81,82^ MEG^83^ and fMRI^84,85^ and, more recently, to determine hierarchical configurations in the brain.^86^ The full mathematical description of the Hopf model is available in the Supplementary material.

To model the whole-brain dynamics requires modelling the coupling of the local dynamics of each brain region interconnected through a given coupling connectivity matrix, *C*. Here, we use a pseudo-gradient procedure to optimize *C*, initially the single-subject structural connectivity matrix, in order to fit empirical data. Specifically, we iteratively compared the output of the model with empirical measures of the FC matrix. Ultimately, GEC is generative, using the whole-brain model to fine-tune the strength of existing anatomical connectivity (e.g., the effective conductive values of each fibre). The full mathematical description of the optimization procedure, its advantages over other models and how we obtained each single-subject structural connectivity, are available in the Supplementary material.

### Areas of interest

Analyses of FC and GEC were organised into three different neurofunctional domains, starting out from primary auditory processing, then radiating out to increasingly deeper levels of processing, first the language network’s syntactic and semantic regions, second the insula, thirdly the hippocampal region. Specifically, our networks and regions of interest were:

(i) The auditory speech processing pathways starting from primary auditory cortex (PAC) and then including planum temporale (PT), posterior superior temporal gyrus (pSTG), planum polare (PP) and anterior superior temporal gyrus (aSTG).^87^ This further included the classical fronto-temporal perisylvian language regions such as the inferior frontal gyrus (IFG), comprising both the pars opercularis (OPER) and pars triangularis (TRI), and superior and middle temporal gyrus (STG and MTG).^88–91^ We also included the semantic network, as meaning is inherent to any act of language use – specifically the anterior temporal lobe (ATL) as a hub of a supramodal concepts associated with words,^92^ and the semantic control network, which is taken to be involved in accessing and manipulating theses stored meaning representations within the contexts they occur in,^45^ specifically right and left posterior inferior frontal gyrus (pars triangularis and pars opercularis; pIFG), the insula, the left posterior middle temporal gyrus (lpMTG), the left posterior inferior temporal gyrus (lpITG), the dorsomedial prefrontal cortex (dmPFC), and the angular gyrus (AG).^47^

(ii) The anterior insula (AI), which plays a crucial role in social communication and ASD,^93,94^ and is a key node in the salience network (SAN), orienting attention towards important stimuli. As such, its dysfunction in ASD may be linked to the fact that people with ASD struggle to find social stimuli salient and meaningful.^49^

(iii) The hippocampus and the regions that comprise it (the tail, body and head), which interacts with language function in the organization of language-mediated memory.^95^

These areas of interest were derived from those regions in the Schaefer atlas that overlap with their corresponding regions in the Yale Brain Atlas (YBA). For example, the IFG in our whole-brain matrix were those regions that overlap with the IFG in the YBA. This is based on the fact that the YBA is region-labelled, whereas the Schaefer atlas is categorized by networks.

### Statistical analysis

Independent two-sample t-tests were computed to detect brain areas displaying significant differences in FC (z-value) patterns between the two groups across all regions mentioned above. P-values under 0.05 were considered indicative of statistical significance. Corrections for multiple comparisons to control for false positives was carried out using the False Discovery Rate (FDR). All significant measures in the nvASD sample as compared to the TD controls were compared to results from the neurotypical control subjects under sedation as compared to the same individuals at rest, in the independent propofol dataset. In this latter sample, paired two-sample t-tests were computed and FDR-corrected. Given the non-parametric distribution of GEC, Mann-Whitney U tests (Wilcoxon Rank-Sum tests) were applied to compare nvASD and control groups, while Wilcoxon Signed-Rank tests were used for within-subject comparison between awake and sedated states. Below, only group differences in the nvASD dataset that were replicated in the independent propofol dataset are reported; for the results of group comparisons of the latter dataset see the Supplementary material.

Since a large number of comparisons between awake and sedated subjects showed no significant differences, we performed post-hoc control analyses to reassure ourselves that, in this dataset, sedation did indeed reduce connectivity within and between brain networks, particularly in the DMN. In contrast, visual and auditory connectivities were preserved. This is consistent with the studies by Naci *et al.*^69^ (which uses the same dataset as the propofol control dataset used in the present paper), and Boveroux *et al.*^42^ Details about these analyses are provided in the Supplementary material (Supplementary Fig. 6, Supplementary Fig. 7 and Supplementary Fig. 8). Note that while we considered using the ComBat harmonization tool for addressing potential differences between dataset from different scanners;^96^ this was not successful due to the small sample size, which made the tool ineffective.

Finally, we performed multiple regression analyses to study the relationships between brain connectivity and behavioural scores, more specifically, the FC and GEC with the ADOS scores. These analyses were carried out for 7 nvASDs, as two out of the nine subjects did not complete the ADOS. In the regression model, the ADOS scores were treated as independent variables, FC and GEC as dependent variables, and age and gender as control variables. P-values were FDR-corrected using Benjamin-Hochberg procedure to account for multiple comparisons. We also extracted the R-squared value, to represent the variance in FC and GEC explained by the ADOS score, age and gender, and the correlation coefficient to evaluate the strength and direction of the relationships between the two variables.

### Data availability

The nvASD dataset that supports the findings of this study are available on request from the corresponding author. Raw data that support the findings on the propofol control dataset are openly available in OpenNeuro at https://doi.org/10.18112/openneuro.ds003171.v2.0.0.

## Results

### Speech and language networks

#### Auditory speech processing network

Figure 2A shows group differences for the auditory speech processing network. Significant decreases in FC were observed in nvASD relative to their matched typically developing peers between PAC, PT, and pSTG, and the aSTG. When separating regions by hemispheres, there was underconnectivity between the left aSTG and the right PAC; between left aSTG and both left and right PT; between left aSTG and right pSTG; and between left aSTG and left PP. Additionally, FC was reduced between the right aSTG with the left PT and left pSTG, as well as between interhemispheric PAC and PT. GEC similarly revealed significant differences between PAC and PT (*P<0.05*), PP (*P<0.01*) and aSTG (*P<0.05*), and between PP and aSTG (*P<0.05*), in both directions. Interhemispheric regions within PAC, PT, PP and aSTG (*P<0.05*) underconnectivities were observed, in addition to multiple regions across the auditory network from the left to the right hemisphere in both directions (distributions are shown in Supplementary Fig. 2). None of these findings were observed in the propofol dataset (see Supplementary Fig. 3).

**Figure 2.**
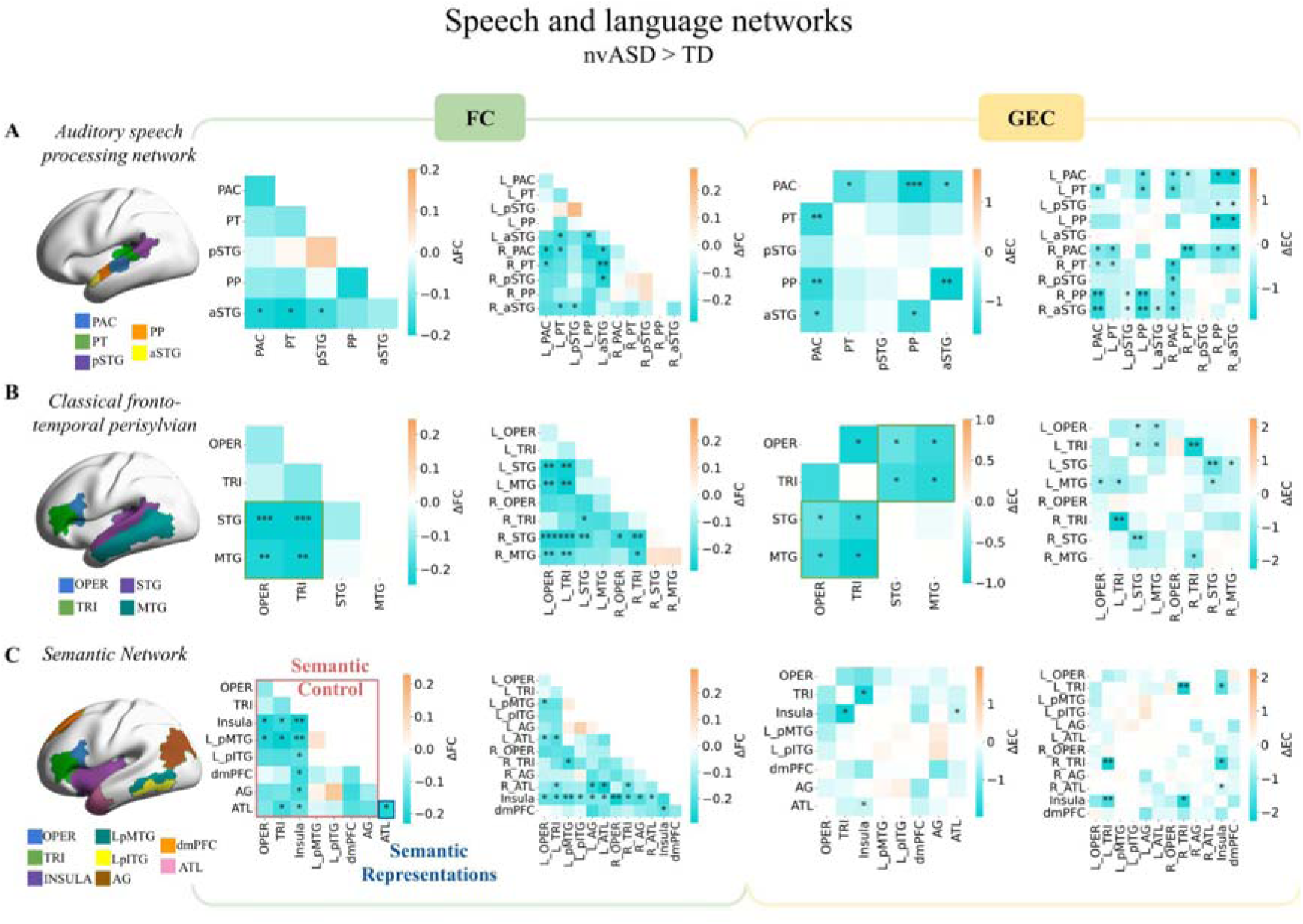
FC and GEC group differences in nvASD compared to controls (ΔFC and ΔGEC). **(A)** the auditory speech processing network, **(B)** Fronto-temporal perisylvian language regions, and **(C)** the semantic network. Bi-hemispheric and hemispherically separated analyses are shown. Diagonal elements in the GEC matrices are zero because we are not considering effective connections within regions, but only between them. Independent two-sample *t*-tests were used for all statistical comparisons. Sample sizes were N = 8 for the typically developing (TD) group and N = 9 for the nvASD group. L_ and R_ prefixes denote regions in the left and right hemispheres, respectively. PAC: primary auditory cortex, PT: planum temporale, pSTG: posterior superior temporal gyrus, PP: planum polare, aSTG: anterior superior temporal gyrus, OPER: operculum, TRI: triangularis, STG: superior temporal gyrus, MTG: middle temporal gyrus, Insula, LpMTG: left posterior middle temporal gyrus, LpITG: left posterior inferior temporal gyrus, dmPFC: dorsomedial prefrontal cortex, AG: angular gyrus, ATL: anterior temporal lobe. Asterisks denote significant differences after FDR correction: * *P<0.05*, ** *P<0.01*, *** *P<0.001*.

#### Classical fronto-temporal perisylvian language network

Figure 2B shows group differences for classical syntactic and semantic language regions. A significant decrease in FC was found in nvASD relative to their controls between both the OPER/TRI and the STG (*P<0.001*) and the MTG (*P<0.01*). These differences corresponded mainly to the left parts of the OPER and TRI, shown in the interhemispheric heatmap. GEC additionally revealed that these underconnectivities may be caused by less connectivity within the left hemisphere: from the left MTG to both left OPER and TRI, and from both left OPER and TRI towards the left STG and left MTG (*P<0.05*). In the right hemisphere underconnectivity was observed from the MTG towards the TRI (*P<0.05*). Additionally, significant interhemispheric underconnectivity was observed in the STG (both in FC, *P<0.01*, and GEC, *P<0.01*) and in TRI (in GEC, *P<0.01*). Intrahemispheric underconnectivity was also observed from the left STG to the right MTG (*P<0.05*), and from the left MTG to the right STG (*P<0.05*) (distributions are shown in Supplementary Fig. 2). These differences again were not observed in the propofol dataset (see Supplementary Fig. 3).

#### Semantic Network

FC differences in nvASD relative to their controls were observed within the semantic network, with a notably reduction in connectivity primarily driven by the insula across the entire network (Figure 2C). As in the perisylvian network above, underconnectivity was detected between the middle temporal region (L_pMTG) and inferior frontal regions (OPER and TRI) (*P<0.05*). Additionally, reduced connectivity was observed both within the ATL and between this region and both the TRI as well as the insula (*P<0.05*). GEC revealed a significant decrease between the insula and both left and right TRI in both directions (*P<0.05*) as well as from the right ATL towards the insula (*P<0.05*) (distributions are shown in Supplementary Fig. 2). The propofol dataset showed significant differences in dmPFC connectivity with L_pITG, AG and ATL in FC, and dmPFC-AG connectivity in GEC, a pattern not observed in the nvASD dataset results (see Supplementary Fig. 3).

### Anterior Insula

Figure 3 shows results for group comparisons of the anterior insula. A strong decrease in FC within the anterior insula was observed (*P<0.01*) as well as an underconnectivity between the anterior insula across both fronto-temporal perisylvian and semantic language regions except for the L_pITG. GEC confirmed effective underconnectivity between the anterior insula with the TRI (*P<0.001*) and the STG (*P<0.05*) in both directions, together with underconnectivities in the fronto-temporal perisylvian network (between the temporal and inferior frontal regions) (distributions are shown in Supplementary Fig. 2). Underconnectivities related to dmPFC in the semantic network are not taken into account as they were also seen to propofol sedation (see Supplementary Fig. 3 and Supplementary Fig. 4).

**Figure 3.**
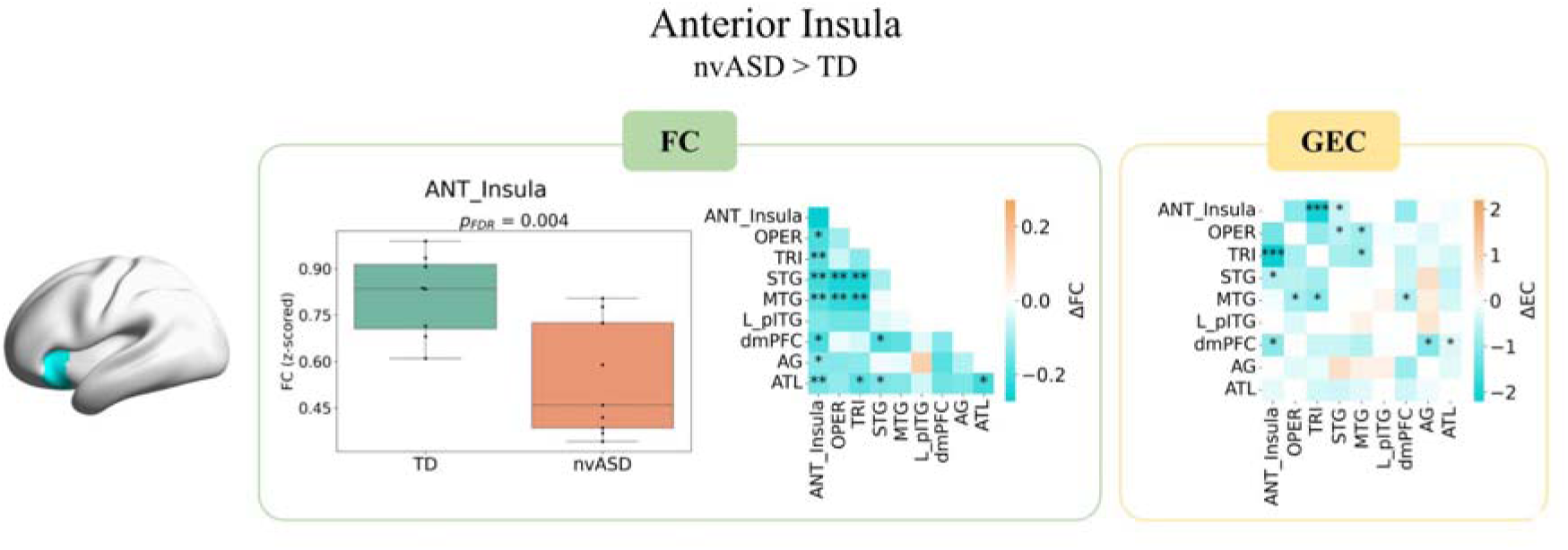
FC and GEC group differences in nvASD vs. controls (ΔFC and ΔGEC). FC inside the anterior insula is illustrated through a boxplot (left); connectivity between the anterior insula and regions from the perisylvian and semantic networks (middle); GEC results for the bi-hemispheric analysis (right). Independent two-sample *t*-tests were used for all statistical comparisons. Sample sizes were N = 8 for the typically developing (TD) group and N = 9 for the nvASD group. ANT_Insula: anterior insula, OPER: operculum, TRI: triangularis, STG: superior temporal gyrus, MTG: middle temporal gyrus, LpITG: left posterior inferior temporal gyrus, dmPFC: dorsomedial prefrontal cortex, AG: angular gyrus, ATL: anterior temporal lobe. Asterisks denote significant differences after FDR correction: * *P<0.05*, ** *P<0.01*, *** *P<0.001*.

### Hippocampus

Figure 4 shows strong significant differences within the hippocampus in the nvASD group (*P<0.001*). GEC measures revealed that this underconnectivity may be caused by a decrease in the causal interaction between the body and head subregions of the hippocampus (distributions are shown in Supplementary Fig. 2). These differences were not observed under propofol sedation (see Supplementary Fig. 5).

**Figure 4.**
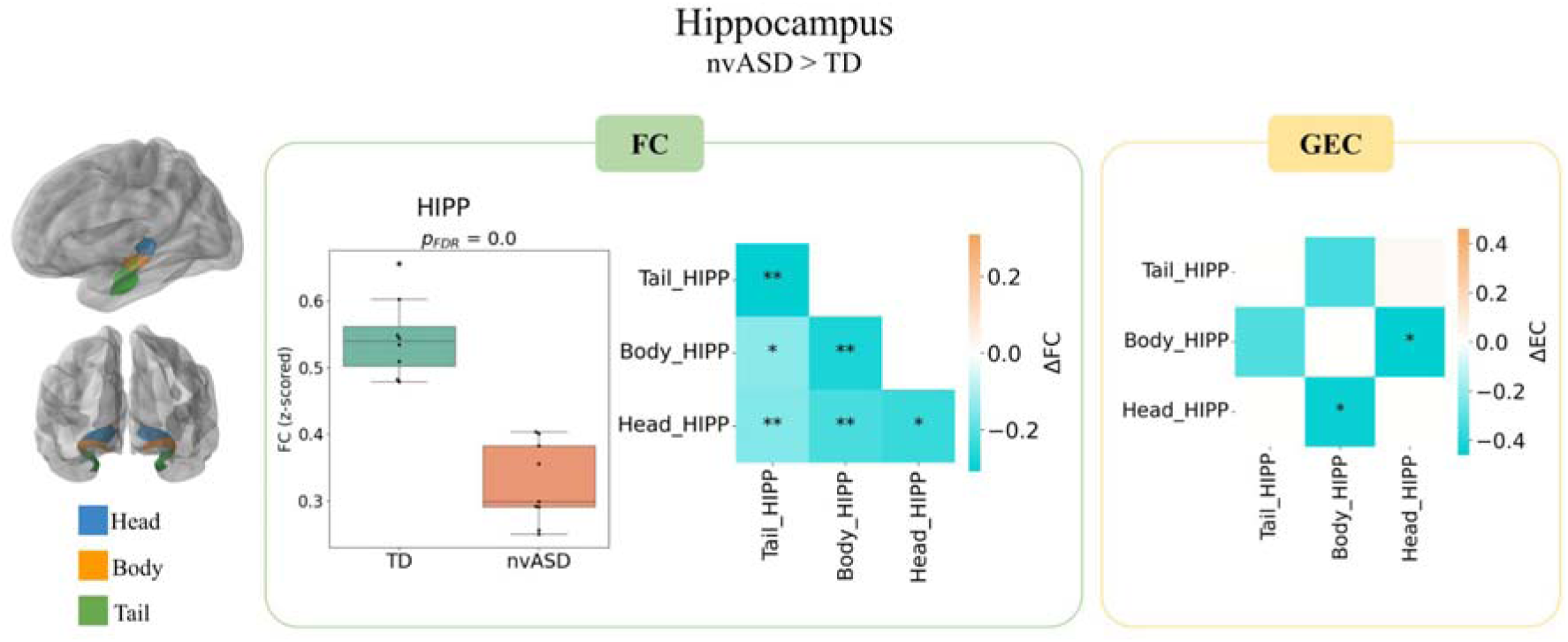
FC and GEC group differences in nvASD vs controls and an independent control dataset of adults scanned sedated and awake (ΔFC and ΔGEC). HIPP: hippocampus. Independent two-sample *t*-tests were used for all statistical comparisons. Sample sizes were N = 8 for the typically developing (TD) group and N = 9 for the nvASD group. Asterisks denote significant differences after FDR correction: * *P<0.05*, ** *P<0.01*, *** *P<0.001*.

### Correlations with ADOS scores

Figure 5 shows significant correlations observed between FC and GEC across all regions analysed above and ADOS total scores, after regressing out the effects of age and gender. In the auditory speech processing network, FC within the left pSTG correlated negatively with the ADOS scores (r=-0.72, *P<0.05*) and its connectivity towards the left PAC correlated strongly and positively with ADOS scores (r=0.73, *P<0.01*). In the frontotemporal perisylvian language network, the FC has negative correlation within the MTG (r=-0.82, *P<0.05*) and the bilateral connection from STG to TRI correlated positively (r=0.94, *P<0.01*). In the semantic network, there was a positive correlation in the FC between the insula and the AG (r=0.91, *P<0.05*) and a negative correlation was obtained between the connectivity of the left insula to left ATL and ADOS (r=-0.95, *P<0.05*).

**Figure 5.**
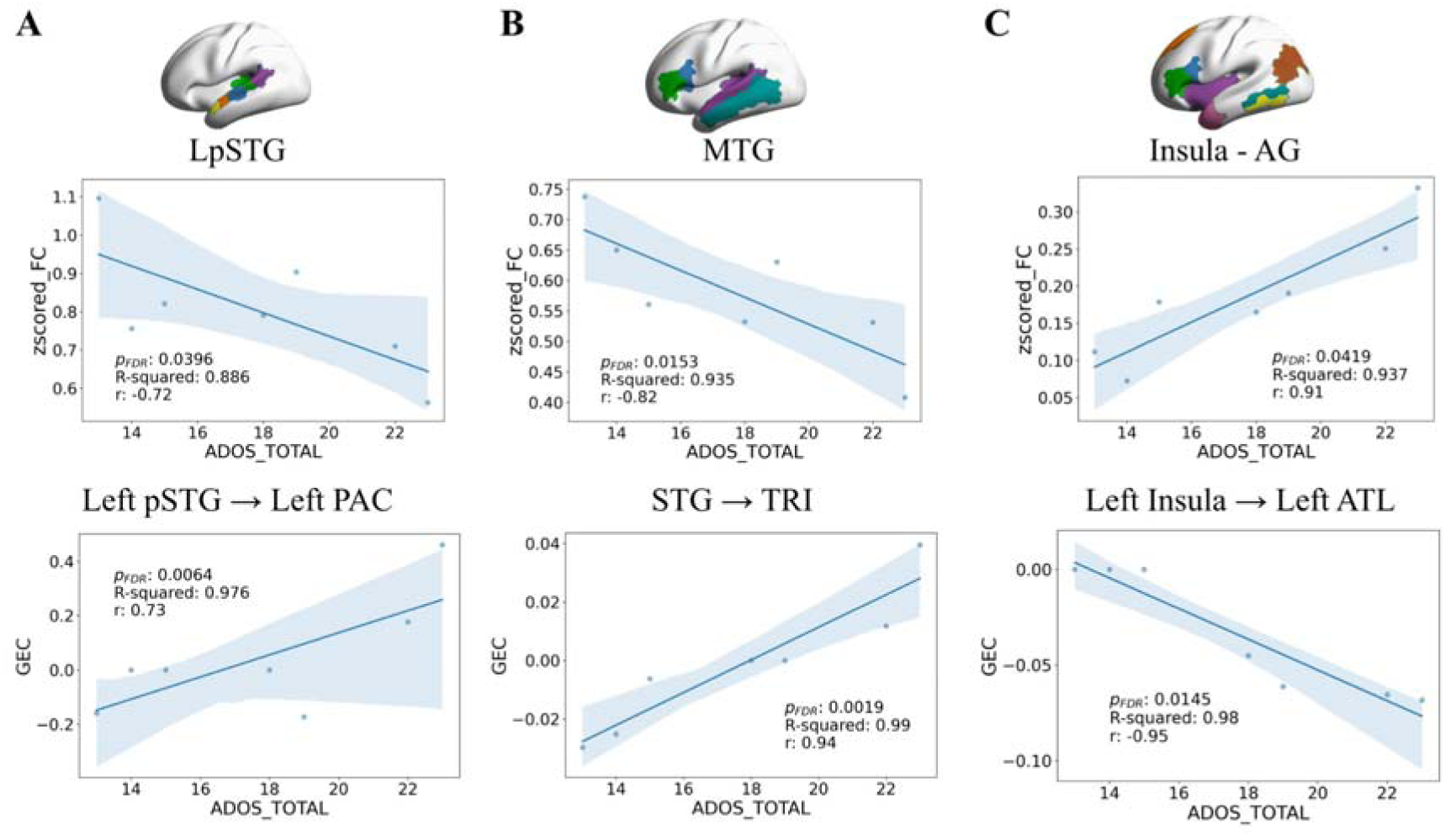
Correlations between ADOS total scores (|r|>0.7, *P<0.05*) and both FC (upper row) and GEC (lower row). **(A)** the auditory speech processing network, **(B)** the frontotemporal perisylvian language network, and **(C)** the semantic network. The statistical test used here is an Ordinary Least Squares (OLS) linear regression. The code fits an OLS model predicting z-scored functional connectivity (dependent variable) from ADOS score (independent variable), while controlling for age and gender. The significance of the ADOS score predictor is assessed using the p-value from the regression, and multiple comparisons are corrected using FDR (False Discovery Rate) correction. Sample sizes were N = 7 for the nvASD group. LpSTG: left posterior superior temporal gyrus, MTG: middle temporal gyrus, AG: angular gyrus. *R-squared* denotes the goodness of fit (closer to 1 represents a better fit) and *r* the correlation between the variables.

## Discussion

This study set out to test for changes in intrinsic brain connectivity in nvASD across a range of neurocognitive networks starting from the auditory language network. Findings show a consistent pattern of hypoconnectivity across all networks and regions tested, using both classical FC and GEC. As the anatomically constrained GEC is designed to provide an explanatory layer for FC, it is unsurprising that our results are consistent with and partially replicate the pattern seen in the FC analysis. While it is impossible to rule out sedation confounds completely, four considerations lend some confidence to the conclusion that absence of language development can be tracked to a widespread loss of synchronization of functional activity in the brain: first, we performed analyses on primary sensory networks (visual and auditory) given prior evidence for preserved connectivity in these networks under propofol sedation;^42^ second, we performed the exact same analyses to an independent dataset of young adults scanned both awake and under propofol, in which no such hypoconnectivity emerged; third, we ran correlational analyses with behavioral ADOS scores within the nvASD group, enhancing interpretability by linking connectivity to the clinical phenotype; and fourth, we conducted control analyses in non-language-related networks. We fully recognize that these steps cannot definitively exclude the possibility that hypoconnectivity patterns are, to some extent, influenced by sedation effects. The only conclusive way to resolve this would have been to scan the nvASD participants both awake and sedated, which, given their level of functioning, remains practically impossible. Notably, beyond these control steps, our findings further indicate that while hypoconnectivity spreads beyond language to key neurocognitive networks involved in salience and semantic processing, and memory–consistent with broad cognitive impairment seen in nvASD–it did not extend to the visual modality in nvASD. Indeed, vision can be a specific strength in ASD at large, and be unaffected even when auditory processing widely is, with increased FC to visual cortices implying worsening language abilities.^97^

A hypoconnectivity pattern was also observed in one of the few fMRI studies of low-functioning (but verbal) children with ASD,^60^ albeit only relative to high-functioning children with ASD. In turn, inter-hemispheric hypoconnectivity was prominent in a rare study of FC (derived from awake fMRI) in a group of nonverbal or low-verbal children across a range of neurogenetic syndromes,^98^ consistent with the present evidence. In high-functioning ASD, a more mixed pattern of local hyperconnectivity within-networks and hypoconnectivity at the level of global network integration, has been a replicated finding.^58,59^ The more homogeneous pattern of pure hypoconnectivity in the present study may thus well be due to the relative homogeneity of the sample itself, as obtained through the language criterion used.

While hypoconnectivity did not extend to the visual network, here analyzed as a control region (Supplementary Figure 6), it encompassed key regions beyond, but interconnected with, speech-processing regions, crucially including the hippocampus. There now is evidence that the hippocampus, far from being confined to enabling episodic memory, mediates the internal creation of structured situation models generally, whether past-directed or not^99^, and plays a critical role in language functions as well.^52,56^ The moment that language is conceptualized as conveying meaning (i.e., a higher-order semantic system), it is also predicted to be necessarily hooked up with the semantic memory, lexical meaning, and semantic control systems. These semantic systems, going beyond the traditional speech processing network, strongly overlap with the DMN,^47^ which Fernandino and Binder^100^ view as coordinating multiple unimodal and multimodal cortical regions so as to result in ‘an integrated simulation of phenomenological experience’, or ‘embodied situation model’. This view is in turn consistent with the perspective of the DMN as sitting on top of a cortical hierarchy from which cognition at large is integrated and orchestrated,^101^ and the perspective of Yeshurun *et al.*^102^, who view the DMN as an ‘active and dynamic “sense-making” network that integrates incoming extrinsic information with prior intrinsic information to form rich, context-dependent models of situations as they unfold over time’, providing narrative structure to our experience in a socially sharable code. In short, if language is necessarily integrated with meaning and our primary code for creating narrative and sharing experience socially, the loss of this system is naturally expected to incur a widespread breakdown of the world models we create neurotypically. From this viewpoint, nvASD presents us with the limiting case of human cognition where the integrating function of language is lost, leading to a change in the apprehension of meaning^25^ and the loss of a coherent narrative of the world.

At a foundational level, this conclusion further informs the debate of whether language dissociates from thought in the human brain. While current versions of a modular view of the language network continue to support such dissociation,^103,104^ the case of nvASD has not been considered in this debate (but see Hinzen et al.^30^). Yet, in humans, it best approximates the critical test case of language being subtracted from the cognitive matrix in humans. In this test case, according to present evidence, we do not see preserved cognition nor evidence for thought resembling the one that is ordinarily expressed in language. While nonverbal IQ, to the extent that it remains measurable, can be in normal ranges in a subsample of the nvASD spectrum, levels of sense-making as measured by nonverbal tests specifically adapted to this population, and comprehension of meaning never reach normal levels.^23,25,28^, Future analyses of FC in nvASD would use graph-theoretical models, at brain-structural and functional levels, to capture whole-brain organizational principles including its small-world organization. In turn, dimensionality-reduction methods such as diffusion embedding might illuminate shifts in the cortical hierarchy as depicted above,^101^ which independent work already established relates to language function, and semantic processing in particular.^105,106^

## Conclusion

This study set out to provide first insights into a disruption of intrinsic functional connectivity in nvASD across language and language-related networks. Results show a pervasive pattern of temporal de-synchronization, which extended to the anatomically constrained generative effective connectivity, encompassing auditory, language, meaning, executive control, and hippocampal networks and their interactions. While language defines nvASD, its absence in this condition is likely to entail a generalized loss of cognitive integration as required for higher-order functions such as building situation models and making sense of the sensory world, which likely depends on language functionality in its human-specific forms. Deficits in interactions involving the anterior insula and hippocampal regions suggest that the effects of nvASD goes beyond language-specific deficits. These results imply, at the neural level, that nvASD condition affects not only language but the general cognitive architecture as a whole.

## Supplementary material

Supplementary material is available at *Brain* online.

