## Supplementary Material for "Brain function in language and associated networks in non- or minimally verbal children"

1. **Image pre-processing with *fMRIPrep*:**

Results included in this manuscript come from preprocessing performed using *fMRIPrep* 21.0.1 (Esteban, Markiewicz, et al., 2018; Esteban, Blair, et al., 2018), which is based on *Nipype* 1.6.1 (Gorgolewski et al., 2011).

The T1-weighted (T1w) image was corrected for intensity non-uniformity (INU) with *N4BiasFieldCorrection* (Tustison et al., 2010), distributed with ANTs 2.3.3 (Avants et al., 2008), and used as T1w-reference throughout the workflow for each participant. The T1w-reference was then skull-stripped with a *Nipype* implementation of the *antsBrainExtraction.sh* workflow (from ANTs), using OASIS30ANTs as the target template. Brain tissue segmentation of cerebrospinal fluid (CSF), white-matter (WM) and grey-matter (GM) was performed on the brain-extracted T1w using *fast* (FSL 6.0.5.1:57b01774) (Zhang, Brady, and Smith, 2001). Brain surfaces were reconstructed using *recon-all* (FreeSurfer 6.0.1) (Dale, Fischl, and Sereno, 1999), and the brain mask estimated previously was refined with a custom variation of the method to reconcile ANTs-derived and FreeSurfer-derived segmentations of the cortical grey-matter of Mindboggle (Klein et al., 2017). Volume-based spatial normalisation to one standard space (MNI152NLin2009cAsym) was performed through nonlinear registration with *antsRegistration* (ANTs 2.3.3), using brain-extracted versions of both T1w reference and the T1w template.

For each of the BOLD runs per subject, the following preprocessing was performed: First, a reference volume and its skull-stripped version were generated using a custom methodology of fMRIPrep. Head-motion parameters with respect to the BOLD reference (transformation matrices, and six corresponding rotation and translation parameters) are estimated before any spatiotemporal filtering using *mcflirt* (FSL 6.0.5.1:57b01774) (Jenkinson et al., 2002). BOLD runs were slice-time corrected to 1.46s (0.5 of slice acquisition range 0s-2.92s) using 3dTshift from AFNI (Cox and Hyde, 1997). The BOLD time-series (including slice-timing correction when applied) were resampled onto their original, native space by applying the transforms to correct for head-motion. These resampled BOLD time-series will be referred to as *preprocessed BOLD in original space*, or just *preprocessed BOLD*. The BOLD reference was then co-registered to the T1w reference using *bbregister* (FreeSurfer) which implements boundary-based registration (Greve and Fischl, 2009). Co-registration was configured with six degrees of freedom. Several confounding time-series were calculated based on the *preprocessed BOLD*: framewise displacement (FD), DVARS and three region-wise global signals. FD was computed using two formulations following Power (absolute sum of relative motions) (Power et al., 2014) and Jenkinson (relative root mean square displacement between affines) (Jenkinson et al., 2002). FD and DVARS are calculated for each functional run, both using their implementations in *Nipype* (following the definitions by Power et al., 2014). The three global signals were extracted within the CSF, the WM, and the whole-brain masks. Additionally, a set of physiological regressors were extracted to allow for component-based noise correction (*CompCor*) (Behzadi et al., 2007). Principal components were estimated after high-pass filtering the *preprocessed BOLD* time-series (using a discrete cosine filter with 128s cut-off) for the two *CompCor* variants: temporal (tCompCor) and anatomical (aCompCor). tCompCor components are then calculated from the top 2% variable voxels within the brain mask. For aCompCor, three probabilistic masks (CSF, WM and combined CSF+WM) are generated in anatomical space. The implementation differs from that of Behzadi et al. in that instead of eroding the masks by 2 pixels on BOLD space, the aCompCor masks are subtracted a mask of pixels that likely contain a volume fraction of GM. This mask is obtained by dilating a GM mask extracted from the FreeSurfer’s *aseg* segmentation, and it ensures components are not extracted from voxels containing a minimal fraction of GM. Finally, these masks are resampled into BOLD space and binarized by thresholding at 0.99 (as in the original implementation). Components are also calculated separately within the WM and CSF masks. For each CompCor decomposition, the *k* components with the largest singular values are retained, such that the retained components’ time series are sufficient to explain 50 percent of variance across the nuisance mask (CSF, WM, combined, or temporal). The remaining components are dropped from consideration. The head-motion estimates calculated in the correction step were also placed within the corresponding confounds file. The confound time series derived from head motion estimates and global signals were expanded with the inclusion of temporal derivatives and quadratic terms for each (Satterthwaite et al., 2013). Frames that exceeded a threshold of 0.5 mm FD or 1.5 standardised DVARS were annotated as motion outliers. All resamplings can be performed with *a single interpolation step* by composing all the pertinent transformations (i.e. head-motion transform matrices, susceptibility distortion correction when available, and co-registrations to anatomical and output spaces). Gridded (volumetric) resamplings were performed using *antsApplyTransforms* (ANTs), configured with Lanczos interpolation to minimise the smoothing effects of other kernels (Lanczos, 1964). Non-gridded (surface) resamplings were performed using *mri_vol2surf* (FreeSurfer).

Many internal operations of *fMRIPrep* use *Nilearn* 0.8.1 (Abraham et al., 2014), mostly within the functional processing workflow. For more details see the fMRIprep website (fMRIPrep, n.d.).

2. **Generative Effective Connectivity (GEC) Methodology**
   2.1. **The Hopf Model**

To model the whole-brain dynamics requires modelling the coupling of the local dynamics of *N* Hopf nodes interconnected through a given coupling matrix, i.e., by adding a diffusive coupling term representing the input received in node *j* from every other node *i*, which is weighted by the corresponding general effective connectivity (GEC), *C_ji_*. This input is modelled using the common difference coupling, which approximates the simplest (linear) part of a general coupling function (Kringelbach et al., 2023). Then, the stochastic coupled nonlinear differential equation gives the states variables:

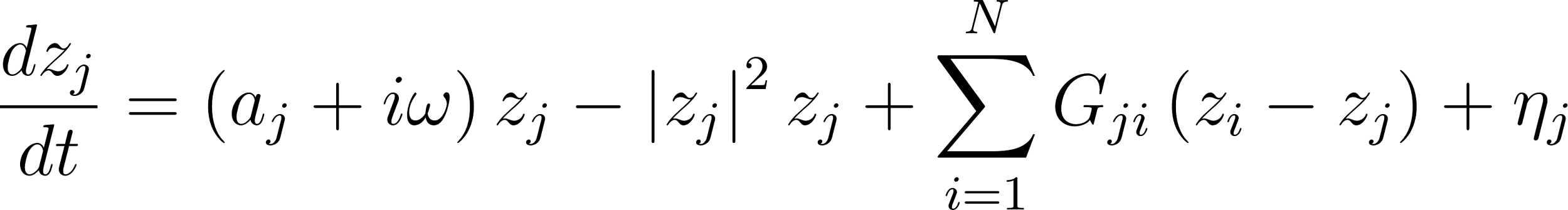
<https://www.codecogs.com/eqnedit.php?latex=%5Cfrac%7Bdz_j%7D%7Bdt%7D%3D%5Cleft%20(%20a_j%2Bi%5Comega%20%5Cright%20)%20z_j-%5Cleft%7C%20z_j%20%5Cright%7C%5E2z_j%20%2B%20%5Csum_%7Bi%3D1%7D%5E%7BN%7DG_%7Bji%7D%5Cleft%20(%20z_i-z_j%20%5Cright%20)%20%2B%5Ceta_j#0>	(1)

where the complex variable (which has arbitrary units)

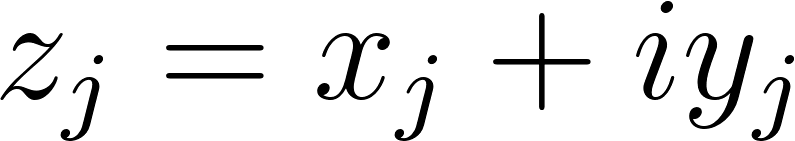
<https://www.codecogs.com/eqnedit.php?latex=z_j%3Dx_j%2Biy_j#0>	(2)

denotes the temporal evolution of the activity in node *j*; |*z_j_*| is the module of *z_j_*, i.e., |*z_j_*|*^2^=x_j_^2^+y_j_^2^*; *ω=2πሀ* is the intrinsic angular frequency (in *rad/s*), where ሀ is the intrinsic frequency in Hz; the parameter *a_j_* is the local bifurcation parameter (in *s^-1^*) and constant across nodes (i.e., *a_j_ =a*); finally, η is the additive white noise, i.e., *〈η(t)〉=0* and *〈η(t)η(t')〉=σ^2^δ(t-t')*, where σ is the noise amplitude (in *s^-1/2^*) fixed to *σ^2^=0.01* (Kringelbach et al., 2023), and the angular brackets *〈.〉* denote the average over stochastic realisations. The intrinsic frequencies ሀ were estimated from empirical data as the averaged peak frequencies of the narrowband BOLD signals of the different nodes (in a 0.008 - 0.08 Hz band, as in Kringelbach et al. 2023).

For $a_{j}<0$, the fluctuations of the local dynamics are attenuated indicating that the system relaxes presenting a stable spiral point producing damped or noisy oscillations in the absence or presence of noise, respectively, in a so-called subcritical regime. On the other hand, if $a_{j}>0$, the system produces self-sustained oscillations, i.e, a stable limit cycle, with a constant amplitude and constant intrinsic frequency $\omega_{j}/(2\pi)$, in a so-called supercritical regime (figure 1S).

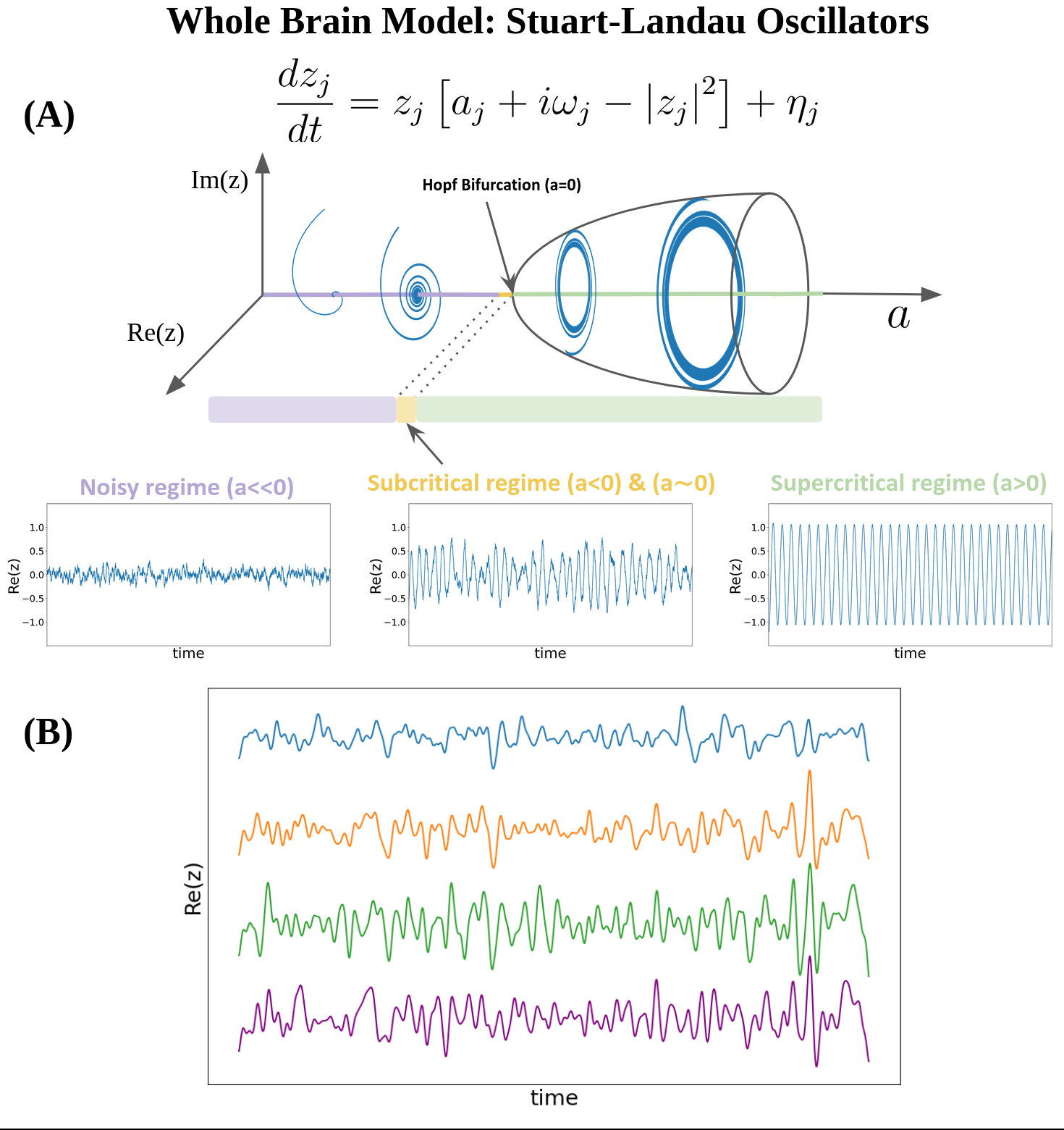

**Supplementary Figure 1**. **(A)** The Stuart-Landau Oscillator produces three different signals given a local bifurcation parameter *a*: 1) Noisy regime - a noise signal resulting from Gaussian noise when the parameter is much less than zero; 2) Subcritical regime - a fluctuating stochastically structured signal when the parameter is just below zero, which allows the system to fluctuate between noise and oscillations; 3) Supercritical regime - an oscillatory signal when the parameter is larger than zero. Parameters: $a_{j}=-0.5, -0.1, 1$; $\omega_{j}=5 rad\cdot s^{-1}$ and $\sigma=0.3$. **(B)** Brain signals are modelled by the real part of the state variables, i.e., *x=Re(x)*. Illustrated are four different brain BOLD signals from the Human Connectome Project (HCP) using Schaefer parcellation with N=100 nodes.

2.2. **Linearization of the Hopf Model**

Because the brain operates near criticality (Chialvo, 2010; Tagliazucchi et al., 2012; Haimovici et al., 2012; Tagliazucchi et al., 2017), the best working point for fitting whole-brain dynamics is at the brink of the Hopf bifurcation, i.e. for the bifurcation parameter *a* at the edge of zero on the negative side, such that the oscillators remain damped still. Previous findings have demonstrated that at this region the correlation between the empirical and the simulated FC is maximised, more specifically, at *a_j_ =-0.02* (Deco et al., 2017; Sanz Perl et al., 2022; Kringelbach et al., 2023).

Previous whole-brain model comparisons showed that non-linearities are minimal. This finding enables us to avoid the nonlinear coupling, thereby simplifying the network statistics without requiring extensive numerical simulations. This suggests that the brain operates in the simpler noisy-oscillation regime and one can estimate the statistics of the whole brain network (e.g, variances and covariances) using a linear approximation (Piccinini et al., 2022; Ponce and Deco, 2024).

The functional correlations between all pairs of brain regions can be estimated by employing a linear noise approximation (LNA), so equation (1) can be re-written as:

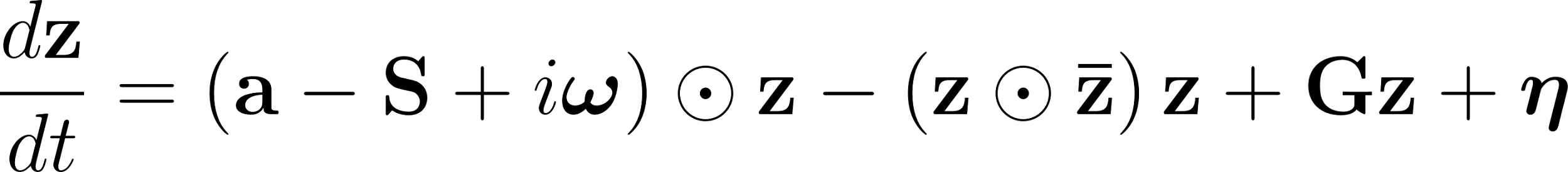
<https://www.codecogs.com/eqnedit.php?latex=%5Cfrac%7Bd%5Cmathbf%7Bz%7D%7D%7Bdt%7D%3D%5Cleft%20(%20%5Cmathbf%7Ba%7D-%5Cmathbf%7BS%7D%2Bi%5Cboldsymbol%7B%5Comega%7D%20%5Cright%20)%5Codot%5Cmathbf%7Bz%7D-%5Cleft%20(%20%5Cmathbf%7Bz%7D%5Codot%5Cmathbf%7B%5Cbar%7Bz%7D%7D%20%5Cright%20)%5Cmathbf%7Bz%7D%2B%5Cmathbf%7BGz%7D%2B%5Cboldsymbol%7B%5Ceta%7D#0>	(1s)

where
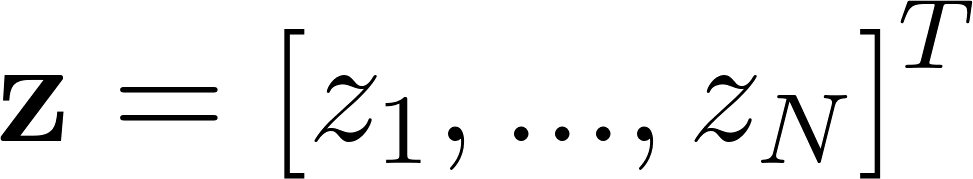
<https://www.codecogs.com/eqnedit.php?latex=%5Cmathbf%7Bz%7D%3D%5Cleft%20%5B%20z_1%2C%20...%2C%20z_N%20%5Cright%20%5D%5ET#0> ,
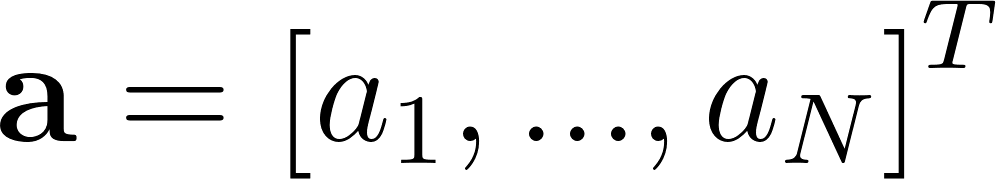
<https://www.codecogs.com/eqnedit.php?latex=%5Cmathbf%7Ba%7D%3D%5Cleft%20%5B%20a_1%2C%20...%2C%20a_N%20%5Cright%20%5D%5ET#0>,
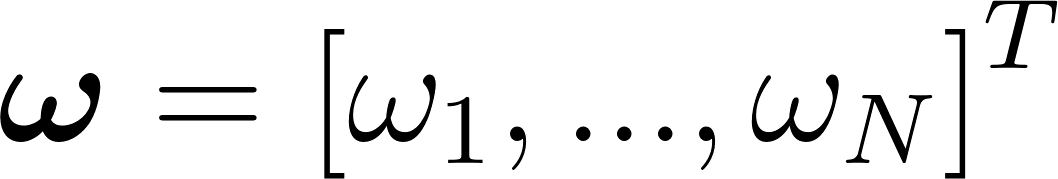
<https://www.codecogs.com/eqnedit.php?latex=%5Cboldsymbol%7B%5Comega%7D%3D%5Cleft%20%5B%20%5Comega_1%2C%20...%2C%20%5Comega_N%20%5Cright%20%5D%5ET#0>,
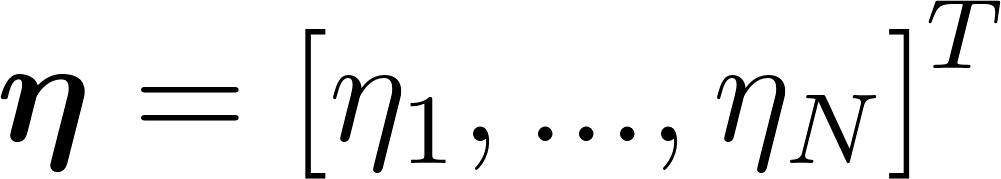
<https://www.codecogs.com/eqnedit.php?latex=%5Cboldsymbol%7B%5Ceta%7D%3D%5Cleft%20%5B%20%5Ceta_1%2C%20...%2C%20%5Ceta_N%20%5Cright%20%5D%5ET#0> represents the vector of uncorrelated noise, and
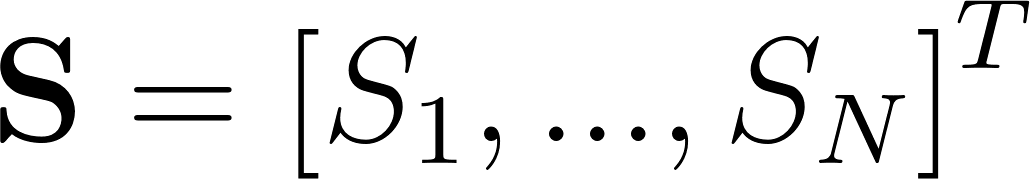
<https://www.codecogs.com/eqnedit.php?latex=%5Cmathbf%7BS%7D%3D%5Cleft%20%5B%20S_1%2C%20...%2C%20S_N%20%5Cright%20%5D%5ET#0> is the vector containing the connectivity strength of each node, i.e, $S_{i}=\Sigma_{j}C_{ij}$ . The superscript *T* represents the transpose,
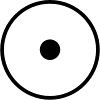
<https://www.codecogs.com/eqnedit.php?latex=%5Codot#0> the Hadamard element-wise product, i.e.,
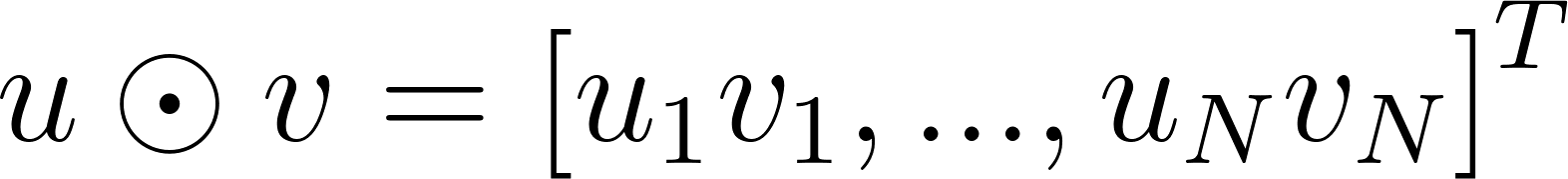
<https://www.codecogs.com/eqnedit.php?latex=u%5Codot%20v%3D%5Cleft%20%5B%20u_1v_1%2C%20...%2Cu_Nv_N%20%5Cright%20%5D%5ET#0>, and
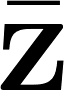
<https://www.codecogs.com/eqnedit.php?latex=%5Cbar%7B%5Cmathbf%7Bz%7D%7D#0> the complex conjugate of
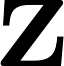
<https://www.codecogs.com/eqnedit.php?latex=%5Cmathbf%7Bz%7D#0>. This equation represents the linear fluctuations
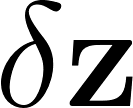
<https://www.codecogs.com/eqnedit.php?latex=%5Cdelta%20%5Cmathbf%7Bz%7D#0> around the fixed point
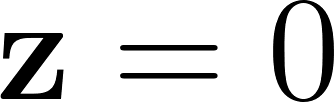
<https://www.codecogs.com/eqnedit.php?latex=%5Cmathbf%7Bz%7D%3D0#0>, which is the solution to the equation
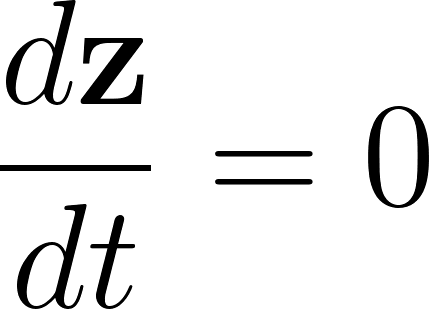
<https://www.codecogs.com/eqnedit.php?latex=%5Cfrac%7Bd%5Cmathbf%7Bz%7D%7D%7Bdt%7D%3D0#0>. Separating the real and imaginary parts, and discarding the higher-order terms
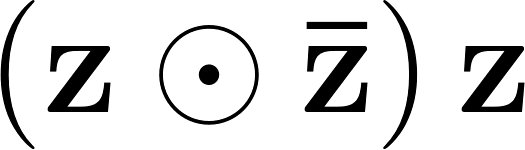
<https://www.codecogs.com/eqnedit.php?latex=%5Cleft(%5Cmathbf%7Bz%7D%5Codot%5Cmathbf%7B%5Cbar%7Bz%7D%7D%5Cright)%5Cmathbf%7Bz%7D#0>, the evolution of the linear fluctuations
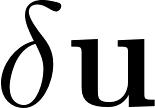
<https://www.codecogs.com/eqnedit.php?latex=%5Cdelta%20%5Cmathbf%7Bu%7D#0> follows a Langevin stochastic linear equation:

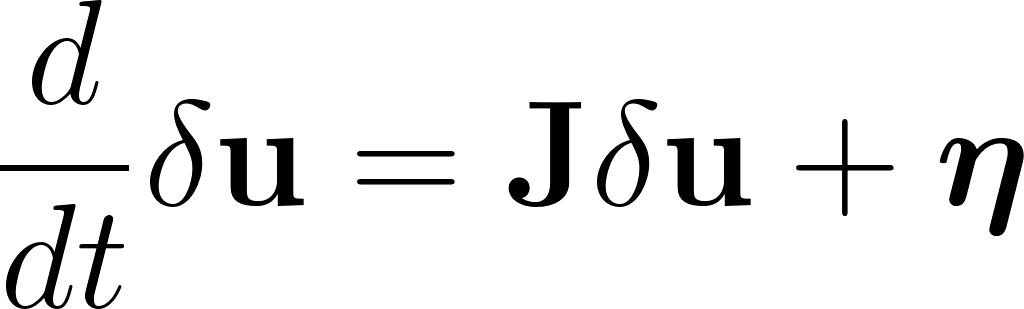
<https://www.codecogs.com/eqnedit.php?latex=%5Cfrac%7Bd%7D%7Bdt%7D%5Cdelta%5Cmathbf%7Bu%7D%3D%5Cmathbf%7BJ%7D%5Cdelta%5Cmathbf%7Bu%7D%2B%5Cboldsymbol%7B%5Ceta%7D#0>,	(2s)

where
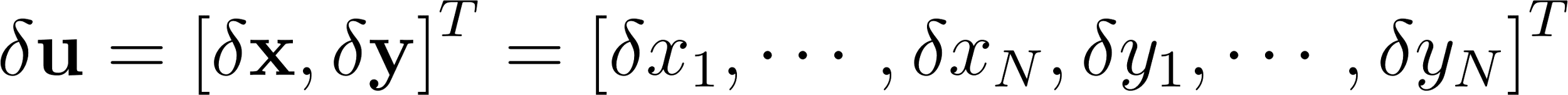
<https://www.codecogs.com/eqnedit.php?latex=%5Cdelta%5Cmathbf%7Bu%7D%3D%5Cleft%5B%20%5Cdelta%5Cmathbf%7Bx%7D%2C%20%5Cdelta%5Cmathbf%7By%7D%5Cright%5D%5ET%3D%5Cleft%5B%20%5Cdelta%20x_1%2C%20%5Ccdots%2C%20%5Cdelta%20x_N%2C%20%5Cdelta%20y_1%2C%20%5Ccdots%2C%20%5Cdelta%20y_N%20%5Cright%5D%5ET#0>is a 2N-dimensional column vector that contains the fluctuations of real and imaginary state variables. The matrix
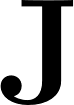
<https://www.codecogs.com/eqnedit.php?latex=%5Cmathbf%7BJ%7D#0> corresponds to the Jacobian of the system evaluated at the fixed point, which can be written as a
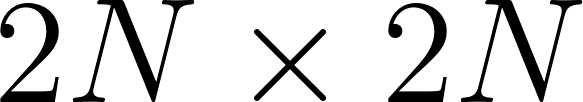
<https://www.codecogs.com/eqnedit.php?latex=2N%5Ctimes%202N#0> matrix

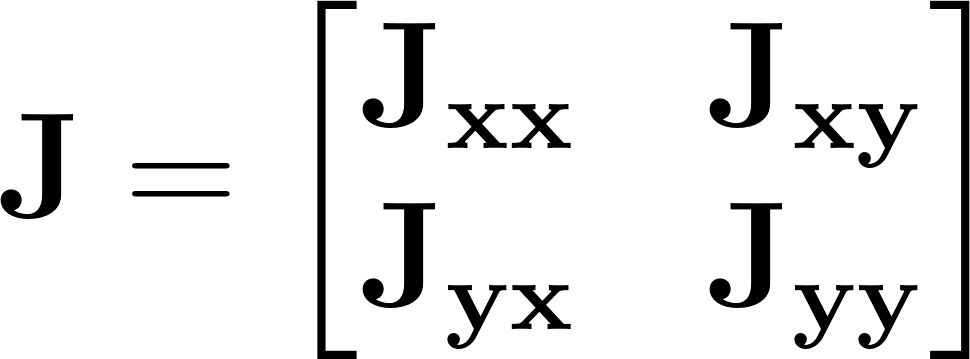
<https://www.codecogs.com/eqnedit.php?latex=%5Cmathbf%7BJ%7D%3D%5Cbegin%7Bbmatrix%7D%20%5Cmathbf%7BJ_%7Bxx%7D%7D%20%26%20%5Cmathbf%7BJ_%7Bxy%7D%7D%20%20%5C%5C%20%5Cmathbf%7BJ_%7Byx%7D%7D%20%26%20%5Cmathbf%7BJ_%7Byy%7D%7D%20%5C%5C%20%5Cend%7Bbmatrix%7D#0>,	(3s)

where
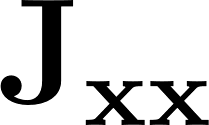
<https://www.codecogs.com/eqnedit.php?latex=%5Cmathbf%7BJ_%7Bxx%7D%7D#0>,
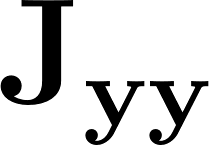
<https://www.codecogs.com/eqnedit.php?latex=%5Cmathbf%7BJ_%7Byy%7D%7D#0>,
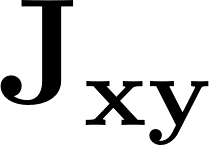
<https://www.codecogs.com/eqnedit.php?latex=%5Cmathbf%7BJ_%7Bxy%7D%7D#0>,
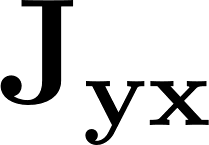
<https://www.codecogs.com/eqnedit.php?latex=%5Cmathbf%7BJ_%7Byx%7D%7D#0>, are
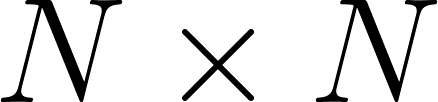
<https://www.codecogs.com/eqnedit.php?latex=N%5Ctimes%20N#0> matrices,
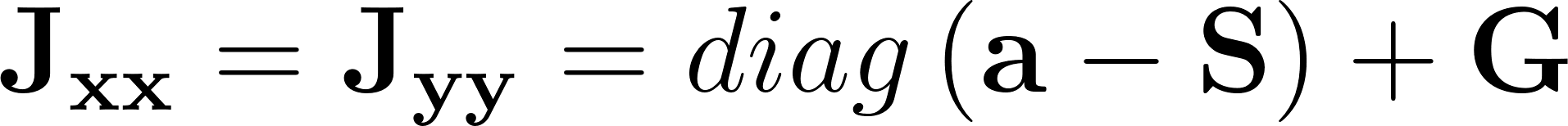
<https://www.codecogs.com/eqnedit.php?latex=%5Cmathbf%7BJ_%7Bxx%7D%7D%3D%5Cmathbf%7BJ_%7Byy%7D%7D%3Ddiag%5Cleft(%20%5Cmathbf%7Ba%7D-%5Cmathbf%7BS%7D%5Cright)%2B%5Cmathbf%7BG%7D#0>, and
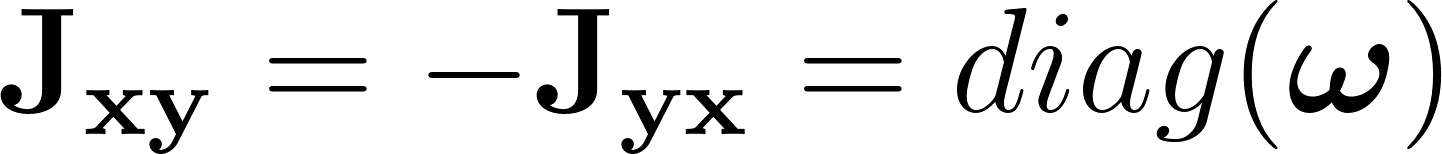
<https://www.codecogs.com/eqnedit.php?latex=%5Cmathbf%7BJ_%7Bxy%7D%7D%3D-%5Cmathbf%7BJ_%7Byx%7D%7D%3Ddiag(%5Cboldsymbol%7B%5Comega%7D)#0>, a diagonal matrix whose diagonal is vector
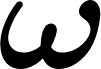
<https://www.codecogs.com/eqnedit.php?latex=%5Cboldsymbol%7B%5Comega%7D#0>. The above linearization is only valid if
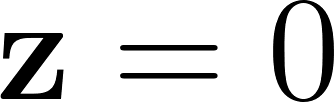
<https://www.codecogs.com/eqnedit.php?latex=%5Cmathbf%7Bz%7D%3D0#0> is a stable solution of the system. Note that all eigenvalues of the Jacobian have a negative real part.

To model the functional connectivity, we must first compute the covariance of the fluctuations around the origin, i.e.,
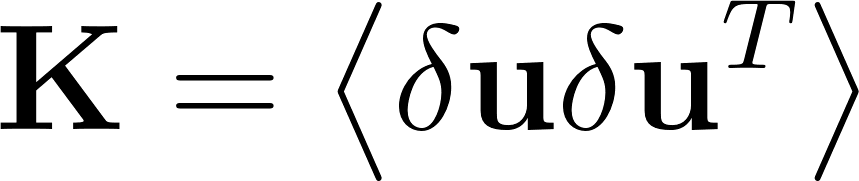
 describing how the fluctuations between regions are related to each other. We begin by writing equation (2s) as
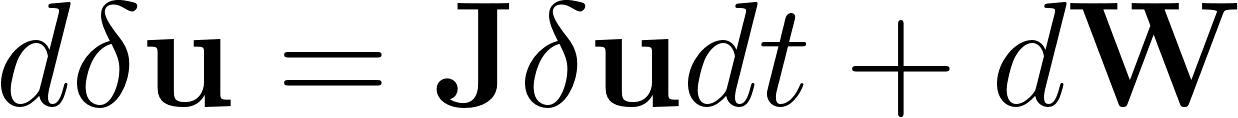
<https://www.codecogs.com/eqnedit.php?latex=d%5Cdelta%5Cmathbf%7Bu%7D%3D%5Cmathbf%7BJ%7D%5Cdelta%5Cmathbf%7Bu%7Ddt%2Bd%5Cmathbf%7BW%7D#0>, where
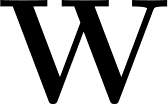
<https://www.codecogs.com/eqnedit.php?latex=%5Cmathbf%7BW%7D#0> is a
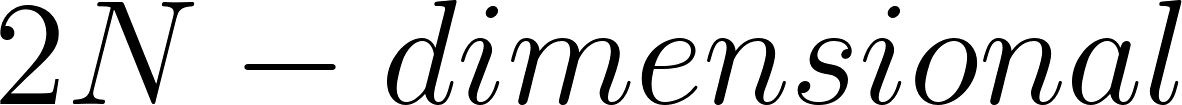
<https://www.codecogs.com/eqnedit.php?latex=2N-dimensional#0> Wiener process (Brownian motion) with covariance
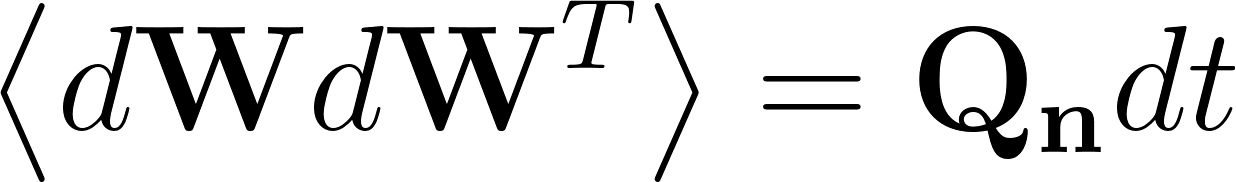
<https://www.codecogs.com/eqnedit.php?latex=%5Cleft%3C%20d%5Cmathbf%7BW%7Dd%5Cmathbf%7BW%7D%5ET%5Cright%3E%3D%5Cmathbf%7BQ_n%7Ddt#0> , and
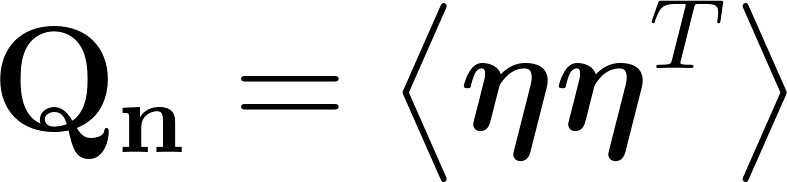
<https://www.codecogs.com/eqnedit.php?latex=%5Cmathbf%7BQ_n%7D%3D%5Cleft%3C%20%5Cboldsymbol%7B%5Ceta%7D%5Cboldsymbol%7B%5Ceta%7D%5ET%20%5Cright%3E#0> is the covariance matrix of the noise (
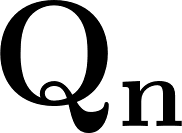
<https://www.codecogs.com/eqnedit.php?latex=%5Cmathbf%7BQ_n%7D#0> is diagonal if the noise is uncorrelated, i.e.,
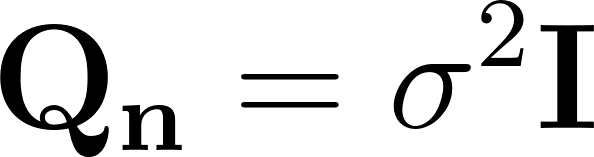
<https://www.codecogs.com/eqnedit.php?latex=%5Cmathbf%7BQ_n%7D%3D%5Csigma%5E2%5Cmathbf%7BI%7D#0>). Then, using Itô’s stochastic calculus, we have
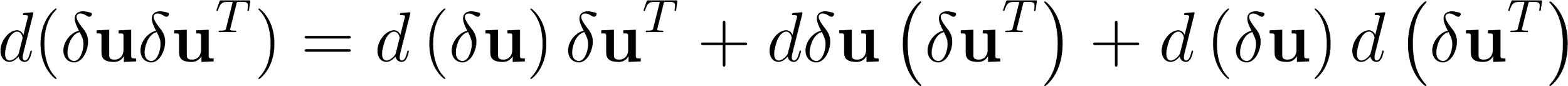
<https://www.codecogs.com/eqnedit.php?latex=d(%5Cdelta%5Cmathbf%7Bu%7D%5Cdelta%5Cmathbf%7Bu%7D%5ET)%3Dd%5Cleft(%5Cdelta%5Cmathbf%7Bu%7D%5Cright)%5Cdelta%5Cmathbf%7Bu%7D%5ET%20%2B%20d%5Cdelta%5Cmathbf%7Bu%7D%5Cleft(%5Cdelta%5Cmathbf%7Bu%7D%5ET%5Cright)%2Bd%5Cleft(%5Cdelta%5Cmathbf%7Bu%7D%5Cright)d%5Cleft(%5Cdelta%5Cmathbf%7Bu%7D%5ET%20%5Cright)#0>. Since
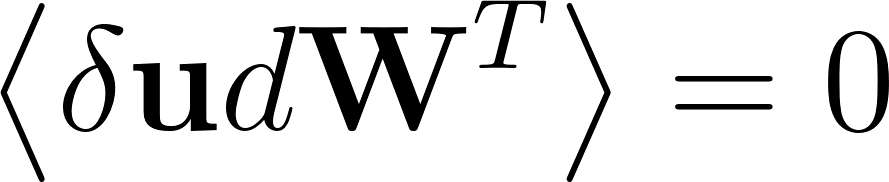
<https://www.codecogs.com/eqnedit.php?latex=%5Cleft%3C%20%5Cdelta%5Cmathbf%7Bu%7Dd%5Cmathbf%7BW%7D%5ET%5Cright%3E%3D0#0> and keeping terms in first order in the differential
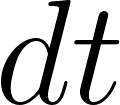
<https://www.codecogs.com/eqnedit.php?latex=dt#0> (as
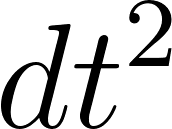
<https://www.codecogs.com/eqnedit.php?latex=dt%5E2#0> can be made arbitrarily small), we obtain
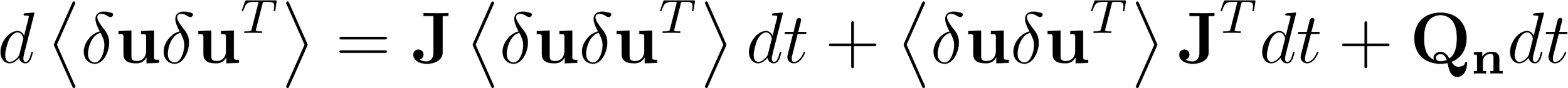
<https://www.codecogs.com/eqnedit.php?latex=d%5Cleft%3C%20%5Cdelta%5Cmathbf%7Bu%7D%5Cdelta%5Cmathbf%7Bu%7D%5ET%5Cright%3E%3D%5Cmathbf%7BJ%7D%5Cleft%3C%20%5Cdelta%5Cmathbf%7Bu%7D%5Cdelta%5Cmathbf%7Bu%7D%5ET%5Cright%3Edt%2B%5Cleft%3C%20%5Cdelta%5Cmathbf%7Bu%7D%5Cdelta%5Cmathbf%7Bu%7D%5ET%5Cright%3E%5Cmathbf%7BJ%7D%5ETdt%2B%5Cmathbf%7BQ_n%7Ddt#0>, and so:

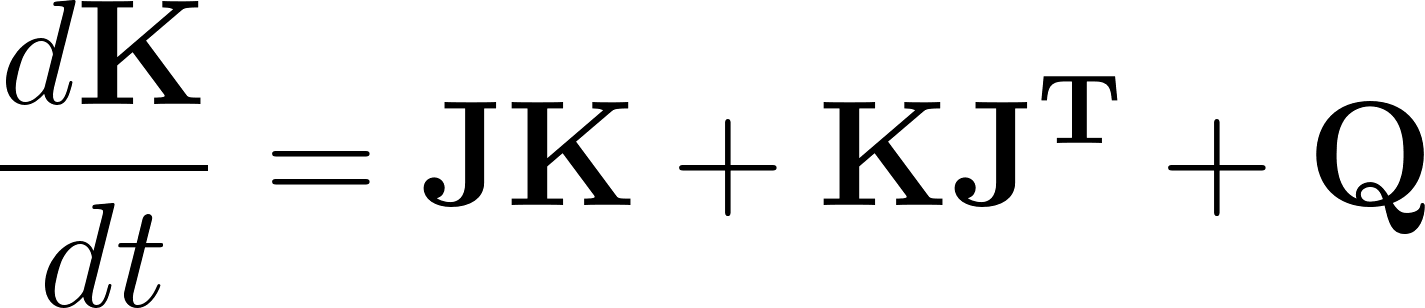
<https://www.codecogs.com/eqnedit.php?latex=%5Cfrac%7Bd%5Cmathbf%7BK%7D%7D%7Bdt%7D%3D%5Cmathbf%7BJK%7D%2B%5Cmathbf%7BKJ%5ET%7D%2B%5Cmathbf%7BQ%7D#0>.	(4s)

Hence, the stationary covariances, for which
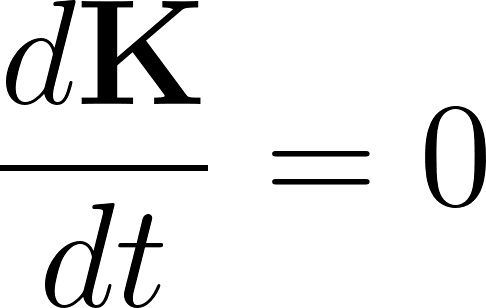
<https://www.codecogs.com/eqnedit.php?latex=%5Cfrac%7Bd%5Cmathbf%7BK%7D%7D%7Bdt%7D%3D0#0>, can be obtained solving the analytic Lyapunov equation using the eigen-decomposition of the Jacobian matrix,

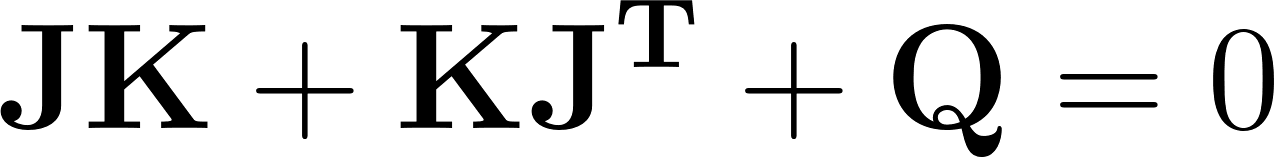
<https://www.codecogs.com/eqnedit.php?latex=%5Cmathbf%7BJK%7D%2B%5Cmathbf%7BKJ%5ET%7D%2B%5Cmathbf%7BQ%7D%3D0#0>.	(5s)

We then obtain the simulated functional connectivity *FC^model^* from the first *N* rows and columns of the covariance
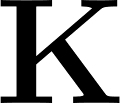
<https://www.codecogs.com/eqnedit.php?latex=%5Cmathbf%7BK%7D#0>, which corresponds to the real part of the dynamics, precisely, representing the BOLD fMRI signal.

2.3. **Model Optimization**

To fit the model to the empirical data, we used a pseudo-gradient procedure to optimize the coupling connectivity matrix *C*, where the starting point was the structural connectivity matrix computed from the diffusion MRI data. The final optimized matrix comprises the effective connectivity values for each pair of anatomical connections instead of the density of fibers. Specifically, we iteratively compared the output of the model with the empirical measures of the functional correlation matrix (*FC^empirical^*), that is, the normalized covariance matrix of the functional neuroimaging data.

We compared the output of the model with *τ* time-shifted covariance (*FS^empirical^(τ)*). These normalized time-shifted covariances were derived by shifting the empirical covariance matrix *KS^empirical^(τ)* and dividing each pair *(i,j)* by $\sqrt{KS_{ii}^{empirical}(0)KS_{jj}^{empirical}(0)}$.

Using a heuristic pseudo-gradient algorithm, we set *α=0.0004* and *ς=0.0001*, and updated *C* until achieving a fully optimized fit, in the form:

$C_{ij}=C_{ij}+\alpha\left( FC_{ij}^{empirical}-{FC}_{ij}^{model} \right) + \varsigma\left( FS_{ij}^{empirical}(\tau)-{FS}_{ij}^{model}(\tau) \right)$,

where *FS^model^(τ)* is defined similarly for *FS^empirical^(τ)*, in other words, it is given by the first *N* rows and columns of the simulated *τ* time-shifted covariance *KS^model^(τ)* normalized by dividing each pair *(i,j)* by $\sqrt{KS_{ii}^{model}(0)KS_{jj}^{model}(0)}$, where *KS^model^(τ)* is the shifted simulated covariance matrix computed as

$KS^{model}(\tau)=exp(\tau J)K$,	(6s)

where *J* is the Jacobian of the linearized Hopf model and *K=KS^model^(0)*. The model is executed iteratively with the updated *C* until a stable and convergent fit is achieved. The update process only modifies the known existing connections from this matrix, following the anatomical connections, and homolog connections between corresponding regions in both hemispheres (as tractography tends to be less accurate in capturing this type of connectivity). In summary, we refer to the optimized matrix *C* as the GEC.

After computing the GEC, normalisation within each individual was performed by the z-score for all existing effective links: (GEC-mean(GEC))/std(GEC). The aim of this is to enable similar contributions of each participant to the statistics. Small effective connectivities will appear as negative due to the removal of the mean value in the z-score function.

2.4. **Model comparison**

This framework is computationally more efficient than those including hemodynamics, such as the Balloon model (Friston et al., 2000) in dynamic causal modelling (DCM) (Friston et al., 2003), Granger causality (Granger, 1980) and transfer entropy (Schreiber, 2000). This is achieved by reducing the degrees of freedom for each brain region, simplifying the complex non-linearity of their activities, in addition to the use of structural data to exclude non-existing connections. Furthermore, this framework facilitates the analysis of larger networks and allows for whole-brain parcellation,, in contrast to the limited number of regions given by DCM to find the network topology.

3. **Single-Subject Structural Connectivity**

To obtain each individual structural connectivity we used the MRtrix pipeline (Tournier et al., 2019) and FSL. After DWI preprocessing, motion and eddy currents corrections followed by brain extraction, the resulting diffusion tensor served to create the whole-brain tractography. To construct a structural brain network using diffusion MRI (dMRI) data, first these images underwent preprocessing including denoising, removing distortion from eddy currents and head motion, and isolating the brain from the skull. Subsequently, the basis functions for each tissue type – white matter, grey matter and cerebrospinal fluid – were estimated using the dhollander function (Dhollander et al., 2016), in order to enable the computation of fibre orientation distributions through spherical deconvolution (Jeurissen et al., 2014). Separately, white matter tissue segmentation was derived from T1-weighted anatomical images, which was then combined with the white matter fibre orientation distributions to generate the streamlines tractography (10 million streamlines per subject) using the Anatomically-Constrained Tractography (ACT) framework (Smith et al., 2012).

In order to create the connectivity matrices, i.e. connectomes, we used our custom atlas that combines the Schaefer atlas (200 parcellation) with subcortical regions (hippocampus and amygdala) from the Yale Brain Atlas. The number of streamlines that begin in one region and end in another determines the degree of connectivity that exists across all regions of interest. It is required to align the diffusion space where the tractogram is formed and the atlas where the regions are specified, applying linear registration. Finally, the tractography data is transformed into a connectivity matrix by mapping the streamlines to the aligned atlas, ensuring to be symmetric with zero self-connections.

4. **Distributions on nvASD dataset results**

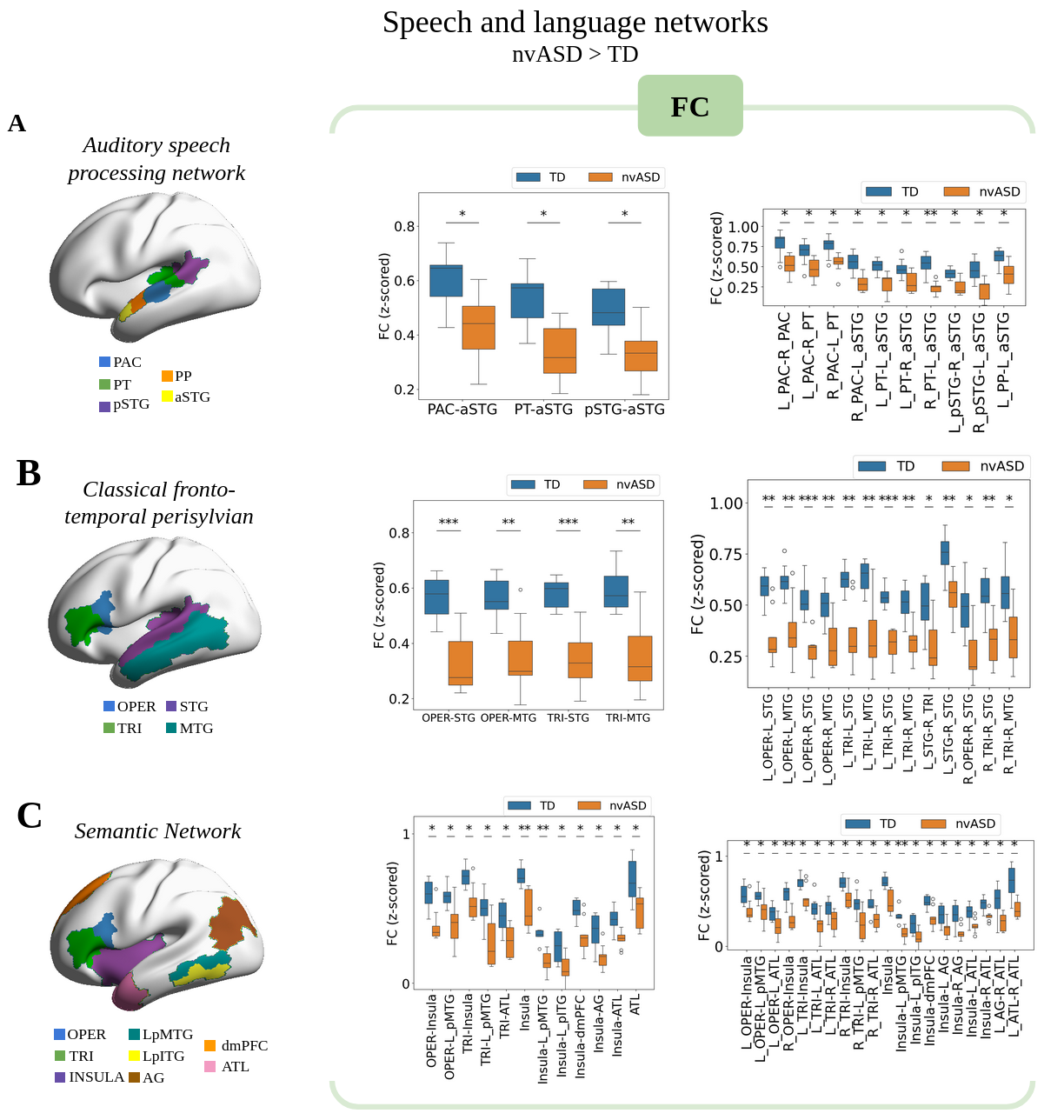

**Supplementary Figure 2A.** Boxplot representation of the distribution for group comparisons across each of the studied networks. Independent two-sample *t*-tests were used for all statistical comparisons. Sample sizes were N = 8 for the typically developing (TD) group and N = 9 for the nvASD group.

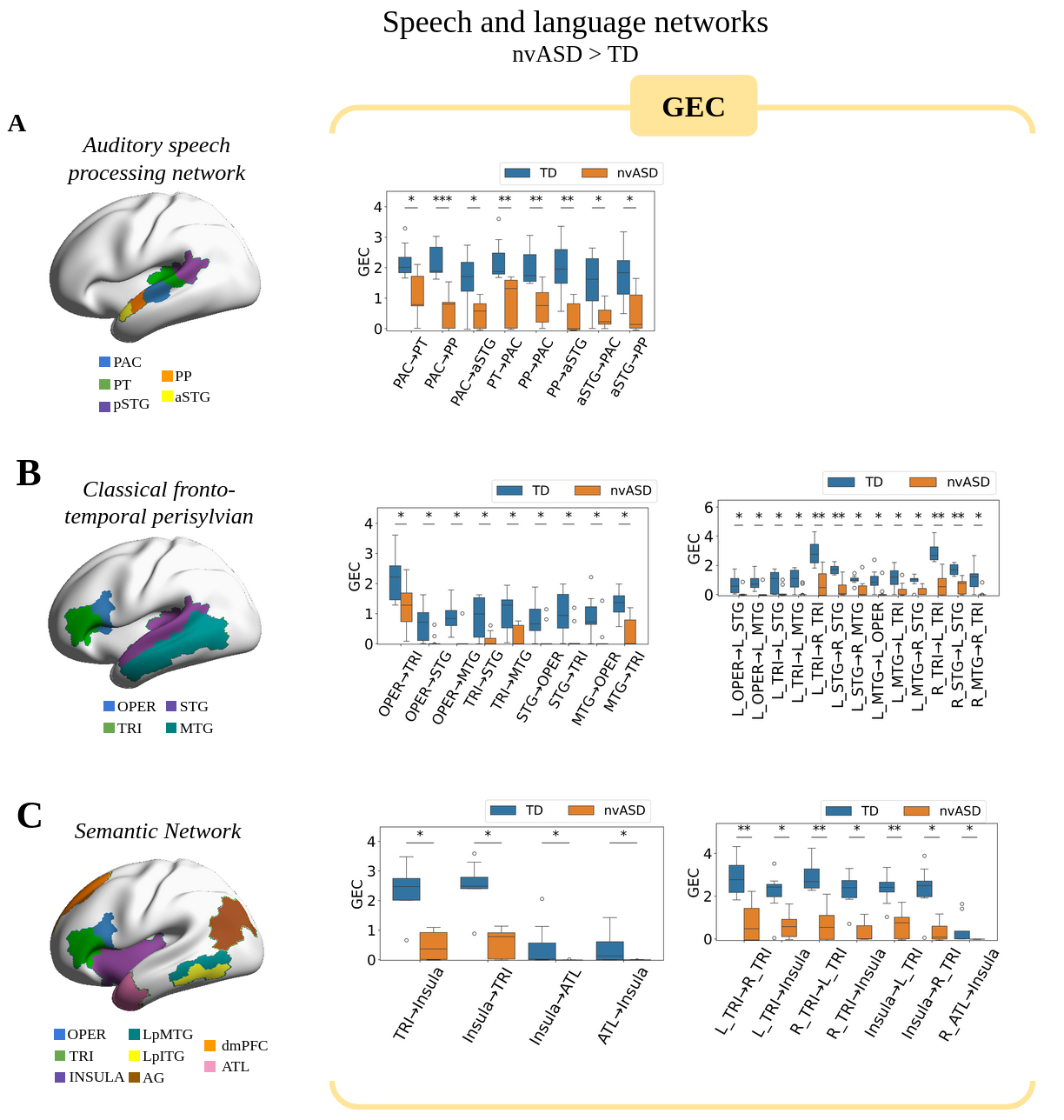

**Supplementary Figure 2B.** Boxplot representation of the distribution for group comparisons across each of the studied networks. Zero values in GEC result from the absence of anatomical connections in the interaction. Independent two-sample *t*-tests were used for all statistical comparisons. Sample sizes were N = 8 for the typically developing (TD) group and N = 9 for the nvASD group.

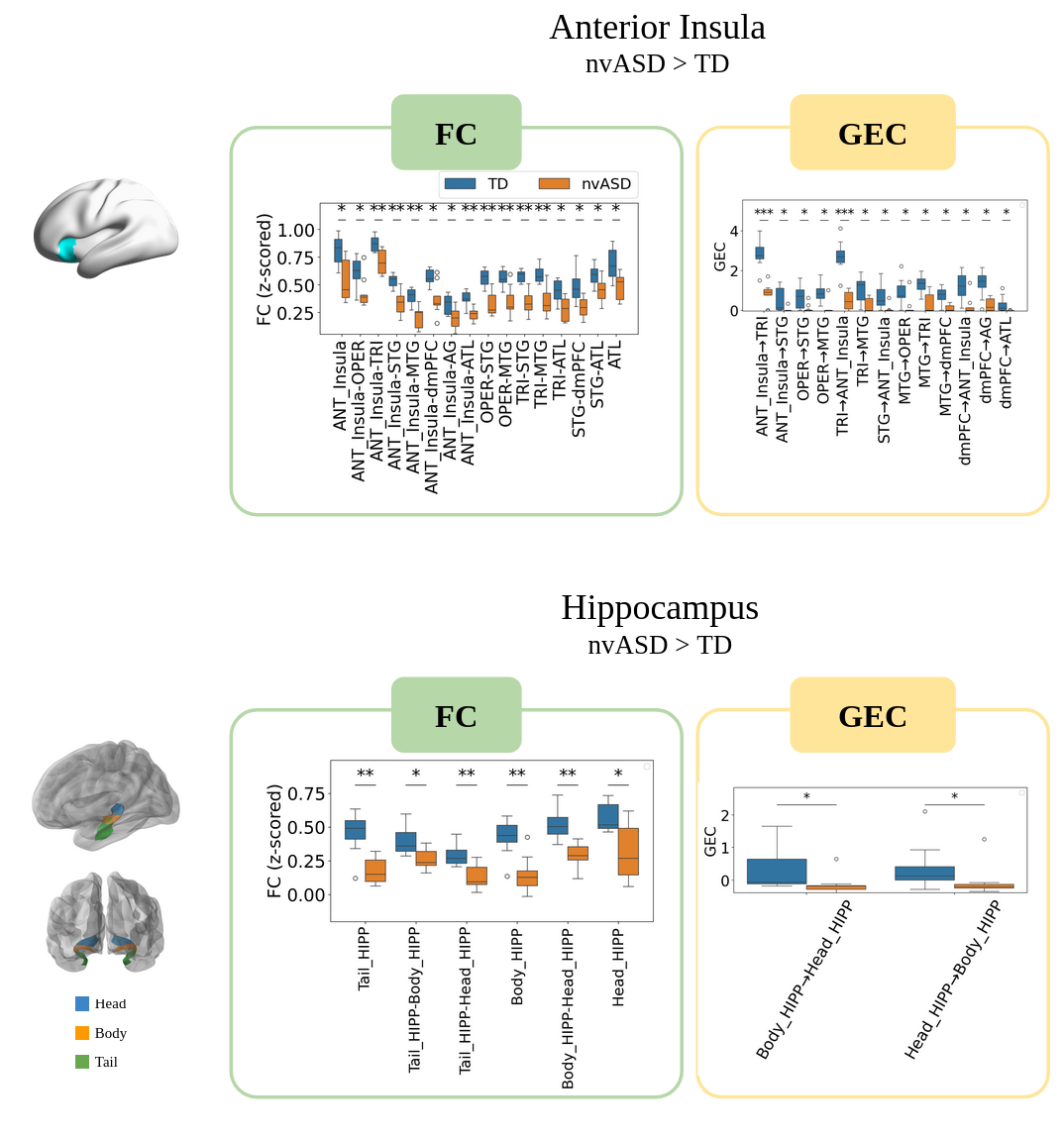

**Supplementary Figure 2C.** Boxplot representation of the distribution for group comparisons across each of the studied networks. Zero values in GEC result from the absence of anatomical connections in the interaction. Independent two-sample *t*-tests were used for all statistical comparisons. Sample sizes were N = 8 for the typically developing (TD) group and N = 9 for the nvASD group.

5. **Results of the independent dataset of neurotypical adults scanned under propofol and awake**
   5.1. **Speech and Language Networks**

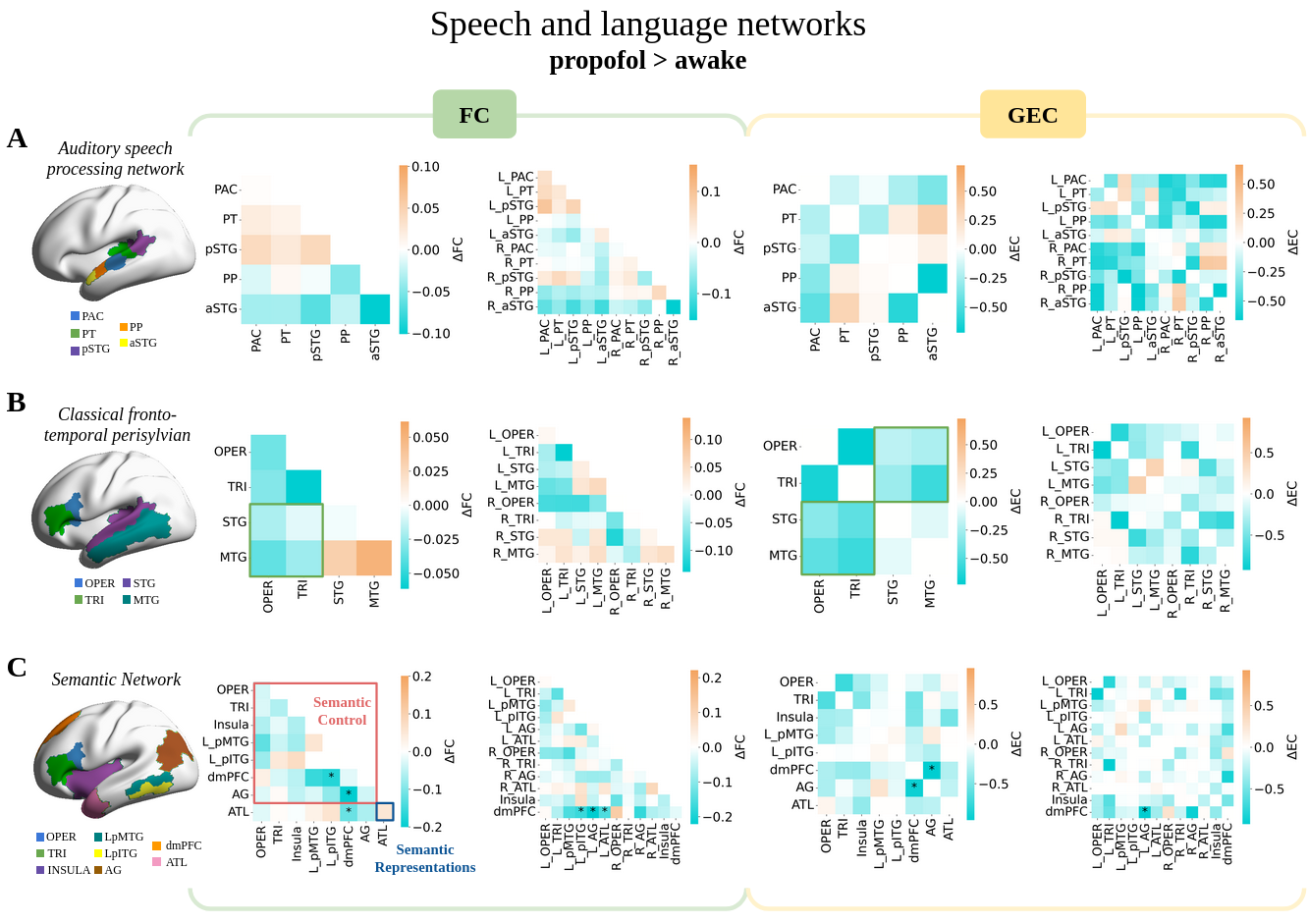
**Supplementary Figure 3**. Dependent two-sample *t*-tests were used for all statistical comparisons. Sample sizes were N = 15. PAC: primary auditory cortex, PT: planum temporale, pSTG: posterior superior temporal gyrus, PP: planum polare, aSTG: anterior superior temporal gyrus, OPER: operculum, TRI: triangularis, STG: superior temporal gyrus, MTG: middle temporal gyrus, Insula, LpMTG: left posterior middle temporal gyrus, LpITG: left posterior inferior temporal gyrus, dmPFC: dorsomedial prefrontal cortex, AG: angular gyrus, ATL: anterior temporal lobe. Asterisks denote significant differences after FDR correction: * *P<0.05*, ** *P<0.01*, *** *P<0.001*.

5.2. **Anterior Insula**

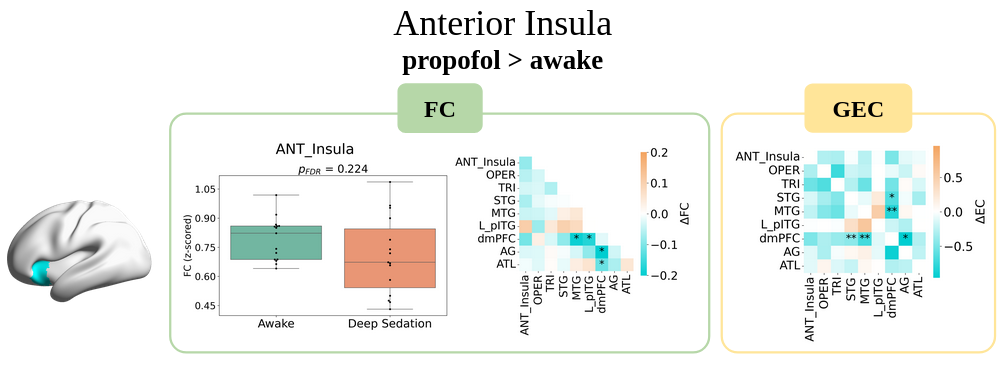
**Supplementary Figure 4**. Dependent two-sample *t*-tests were used for all statistical comparisons. Sample sizes were N = 15. ANT_Insula: anterior insula, OPER: operculum, TRI: triangularis, STG: superior temporal gyrus, MTG: middle temporal gyrus, LpITG: left posterior inferior temporal gyrus, dmPFC: dorsomedial prefrontal cortex, AG: angular gyrus, ATL: anterior temporal lobe. Asterisks denote significant differences after FDR correction: * *P<0.05*, ** *P<0.01*.

5.3. **Hippocampus**

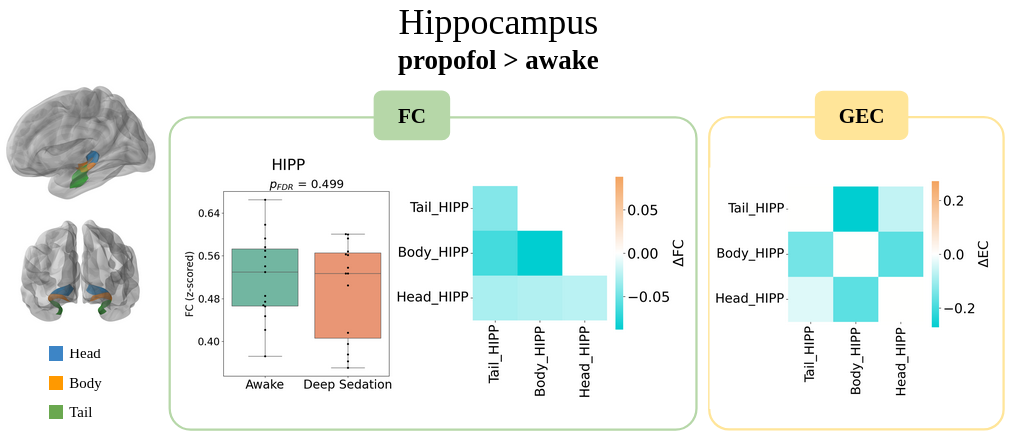

**Supplementary Figure 5**. Dependent two-sample *t*-tests were used for all statistical comparisons. Sample sizes were N = 15. HIPP: hippocampus.

6. **Visual network as a control**

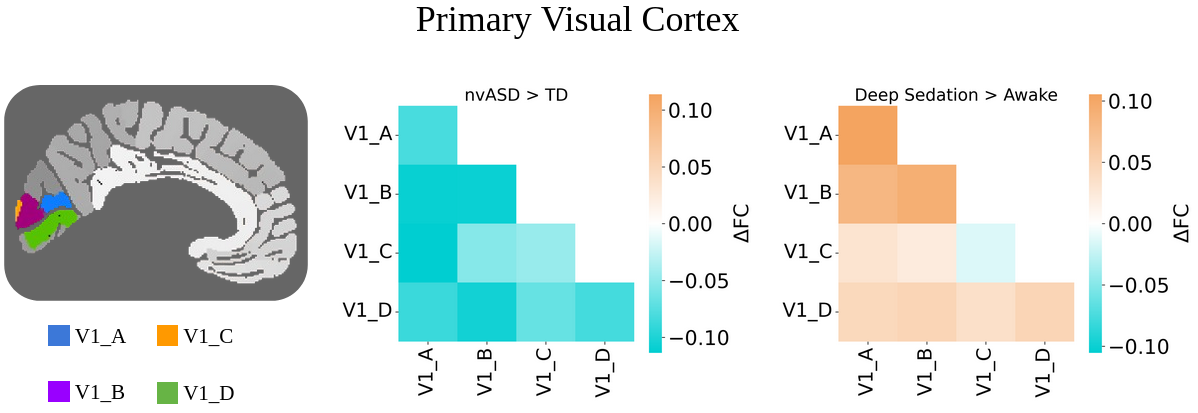
**Supplementary Figure 6**. FC group differences (ΔFC) in nvASD and propofol sedation within regions in the primary visual cortex. Independent two-sample *t*-tests were used for nvASD and HC statistical comparisons, and dependent two-sample *t*-tests were used for awake and propofol statistical comparisons. Sample sizes were N = 8 for the typically developing (TD) group and N = 9 for the nvASD group, and N=15 for the sedated group. V1_A, V1_B, V1_C and V1_D correspond to subregions of the primary visual cortex.

7. **Replication studies in propofol sedation**

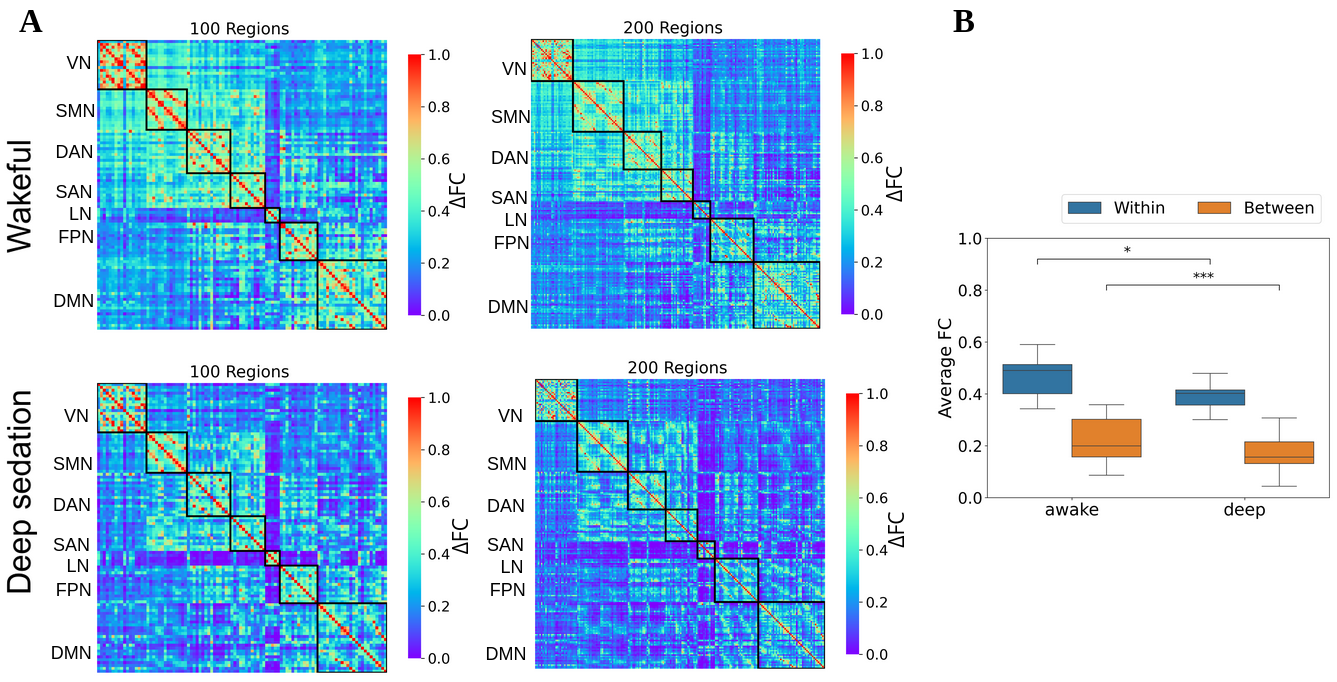
**Supplementary Figure 7**. **(A)** Functional connectivity patterns in seven brain networks in wakeful and deep anaesthesia states for two brain parcellations (Schaefer 100 and 200). A replication study from Naci et al. (2018). **(B)** Our study reveals a significant decrease in FC between and within brain networks. Dependent two-sample *t*-tests were used for all statistical comparisons. Sample sizes were N = 15. VN: Visual Network, SMN: Somatomotor Network, DAN: Dorsal-Attention Network, SAN: Salience Network, LN: Limbic Network, FPN: Frontoparietal Network, DMN: Default Mode Network. ECN: Executive Control Network, AUD: Auditory Network. Asterisks denote significant differences after FDR correction: * *P<0.05*, *** *P<0.001*.

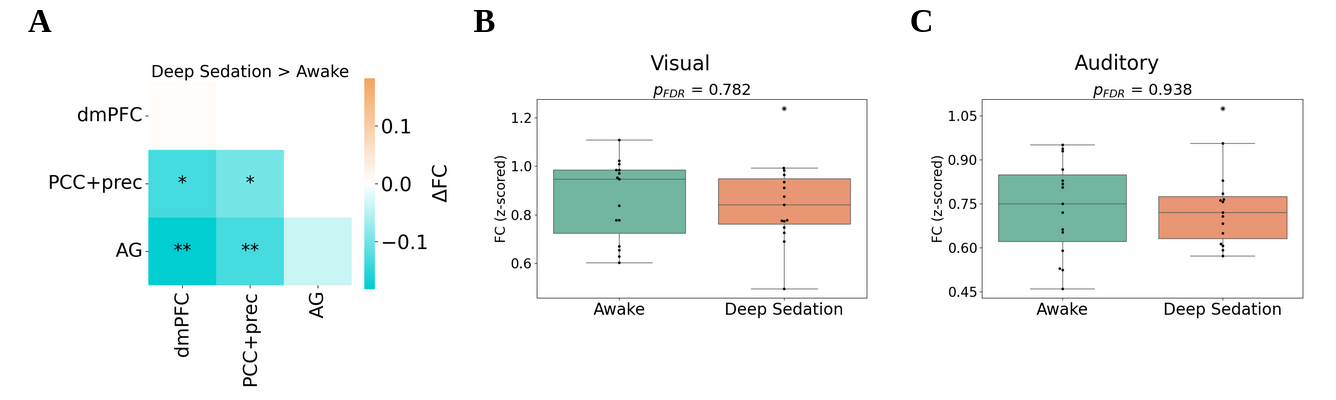
**Supplementary Figure 8**. Our seed-based FC mimics the Bouveroux et al. (2010) –a FC study using ICA in wakeful and deep anaesthesia states– showing **(A)** decreased FC within the DMN and non-significant effects in **(B)** visual and **(C)** auditory regions. Dependent two-sample *t*-tests were used for all statistical comparisons. Sample sizes were N = 15. dmPFC: dorsomedial prefrontal cortex, PCC: posterior cingulate cortex, prec: precuneus, AG: angular gyrus. Asterisks denote significant differences after FDR correction: * *P<0.05*, ** *P<0.01*.
